# Pathogen effector forms a hexameric phosphatase holoenzyme with host core enzyme to promote disease

**DOI:** 10.1101/2025.06.14.659724

**Authors:** Jinlong Wang, Hui Li, Wei Sun, Jiuyu Wang, Xiaodong Fang, Xiaoqi Yang, Chao Liu, Gang Sheng, Wenbo Ma, Yanli Wang

## Abstract

Pathogen effectors play critical roles in pathogenesis by directly manipulating host cellular processes. The Ser/Thr protein phosphatase 2A (PP2A) is commonly targeted by pathogens. Effectors from devastating plant pathogen *Phytophthora* hijack the PP2A core enzyme in plant hosts, altering the host phosphoproteome. In this study, we present a series of cryo-electron microscopy structures of the *Phytophthora* effector PSR2 in complex with the host PP2A core enzyme to form a functional holoenzyme. The PSR2-PP2A complex adopts a unique hexameric architecture, driven by PSR2-mediated dimerization of two heterotrimers. This hexamer exhibits greater stability and features a more exposed catalytic pocket compared to the canonical trimeric form of PP2A, likely enhancing its virulence activity. Mutational analyses underscore the importance of this hexameric structure for PSR2’s virulence function. These findings provide mechanistic insights into pathogen-mediated manipulation of a key host phosphatase and offer targets for developing disease resistance strategies.

## Main

Hosts and pathogens are engaged in a constant co-evolutionary arms race^1^. While plant and animal hosts have evolved sophisticated defense networks to prevent infection by potential pathogens, pathogens employ virulence proteins to undermine host immunity and cellular processes to drive disease progression^2–4^. Understanding the underlying mechanisms of pathogen virulence functions is instrumental in establishing fundamental principles in pathogenesis and developing resistance strategies.

Protein phosphorylation is a key regulatory mechanism in cellular function. The evolutionarily conserved Ser/Thr protein phosphatase 2A (PP2A) is responsible for the majority of dephosphorylation activity occurs on Serine and Threonine residues, thus playing a critical housekeeping role in regulating numerous cellular processes^5–9^. Therefore, dysregulation of PP2A has been found to associate with various diseases^10–12^. PP2A functions as a heterotrimeric holoenzyme composed of a scaffold subunit (A), a catalytic subunit (C) and a regulatory subunit (B). The A and C subunits form the core enzyme, while the B subunit defines substrate specificity, determining which cellular targets are dephosphorylated^13–15^.

Consistent with its important functions, PP2A is a common target for pathogenic manipulation in both plant and animal hosts. For instance, the DNA tumor virus SV40 disrupts human PP2A function through a small antigen, which binds to the PP2A core enzyme, leading to cellular transformation^16–18^. Proteins encoded by human papillomaviruses (HPV) and Epstein-Barr virus (EBV) are also known to associate with the PP2A core enzyme^19,20^. Similar to animal pathogens, plant pathogens also target host PP2A to promote infection^21–23^. In particular, several virulence proteins produced by the devastating *Phytophthora* pathogens have recently been shown to hijack the plant PP2A core enzyme, enhancing infection by altering host phosphoproteome^23^.

*Phytophthora* species, such as the Irish potato famine pathogen *Phytophthora infestans*, are filamentous eukaryotic pathogens that pose significant threats to global plant health and food security^24^. Each *Phytophthora* genome encodes hundreds of virulence proteins called effectors, which possess diverse host-manipulating activities^25,26^. A large family of *Phytophthora* effectors are comprised of tandem repeats of the (L)WY motif, named by conserved residues including multiple leucine (L), one tryptophan (W), and one tyrosine (Y)^27^. The W-Y interaction stabilizes a nearly identical α-helical bundle formed by each (L)WY unit, whereas the leucine residues facilitate stable concatenations through a conserved mechanism^27^. The best-studied (L)WY effector is the *Phytophthora* suppressors of RNA silencing 2 (PSR2), which contains one N-terminal WY unit (named WY1) followed by six LWY motifs (named LWY2 – LWY7). This architecture was named WY1-(LWY)_6_^27^ Importantly, the WY1-(LWY)_n_ arrangement is prevalent in *Phytophthora* species, indicating a role in driving effector evolution. Despite the structural conservation, the (L)WY units are variable in sequences, potentially mediating interactions with different host targets inside the plant cells during infection^27^.

We recently found that 13 WY1-(LWY)_n_ effectors, including PSR2, share a functional module that is based on a specific (L)WY-LWY combination that mediates direct interaction with the A subunit of plant PP2A. These effectors efficiently recruit the host PP2A core enzyme and manipulate the host phosphoproteome as a key virulence mechanism in *Phytophthora* pathogens^23^. By characterizing the crystal structure of PSR2 in complex with a plant PP2A A subunit, we discovered that the PP2A-interacting module clamps onto an N-terminal region of the PP2A A subunit, a region implicated in the interactions with PP2A B subunits in human holoenzymes. Therefore, these PP2A-interacting *Phytophthora* effectors may recruit the PP2A core enzyme as innovative PP2A B subunits^23^. However, the molecular details of the putative effector-PP2A holoenzymes was unknown.

Here, we determined the cryo-electron microscopy (cryo-EM) structure of the PSR2-PP2A holoenzyme complex. We found that the PSR2-PP2A holoenzyme forms a unique hexameric architecture, unlike the canonical hetertrimeric structure of known PP2A holoenzymes. Detailed structural and functional analyses suggest that the hexamerization enables the formation of a more robust holoenzyme, thus enhancing the virulent activity of PSR2. This study provides a deeper understanding of how pathogens manipulate their hosts. As key susceptibility factor across the kingdom, this finding offers new avenues to improve disease resistance through precise engineering of PP2A.

## Results

### PSR2-PDF1-Cα holoenzyme forms a hexamer

To understand how PSR2 hijacks the PP2A core enzyme, we determined the structure of the PSR2-PP2A holoenzyme. *A. thaliana* encodes three PP2A A subunits (PDF1, PDF2 and RCN1) and five C subunits (PP2A-1-PP2A-5)^28^. Since all three A subunits directly interact with PSR2, we selected PDF1 for structural studies. However, none of the five C subunits produced soluble proteins using various expression systems. Therefore, we used the human PP2A C subunit (Cα), which shares a high degree of sequence similarity with its *A. thaliana* counterparts to form a core enzyme with PDF1^23^. Structural predictions using AlphaFold3 indicate a strong conservation between Cα and the *A. thaliana* C subunits, with root-mean-square deviation (RMSD) values ranging from 0.5 to 0.7 Å (Supplementary Fig.1a)^29^. The sequence and structural similarities between the plant and host PP2A C subunits suggest that the PSR2-PDF1-Cα complex likely mimics the native PSR2-PP2A holoenzyme in plant cells.

We assembled the PSR2-PDF1-Cα holoenzyme by incubating purified PDF1 and Cα to form the core enzyme, followed by the addition of purified PSR2. The complex was purified via gel filtration chromatography (Supplementary Fig.1b). We determined the cryo-EM structure of the PSR2-PDF1-Cα complex at an overall resolution of 3.38 Å (Supplementary Fig.2a-2c and Table 1). Interestingly, we found that the PSR2-PDF1-Cα complex forms a hexameric structure, containing two copies of each protein (Fig. 1a). Local resolution distribution revealed that the N-terminus of both PDF1 and PSR2 are flexible, as indicated by lower resolutions in these regions. In contrast, the C-termini of PDF1 and PSR2 exhibited higher stability and better electronic density, allowing for generating detailed atomic models (Supplementary Fig.2d).

**Fig.1.**
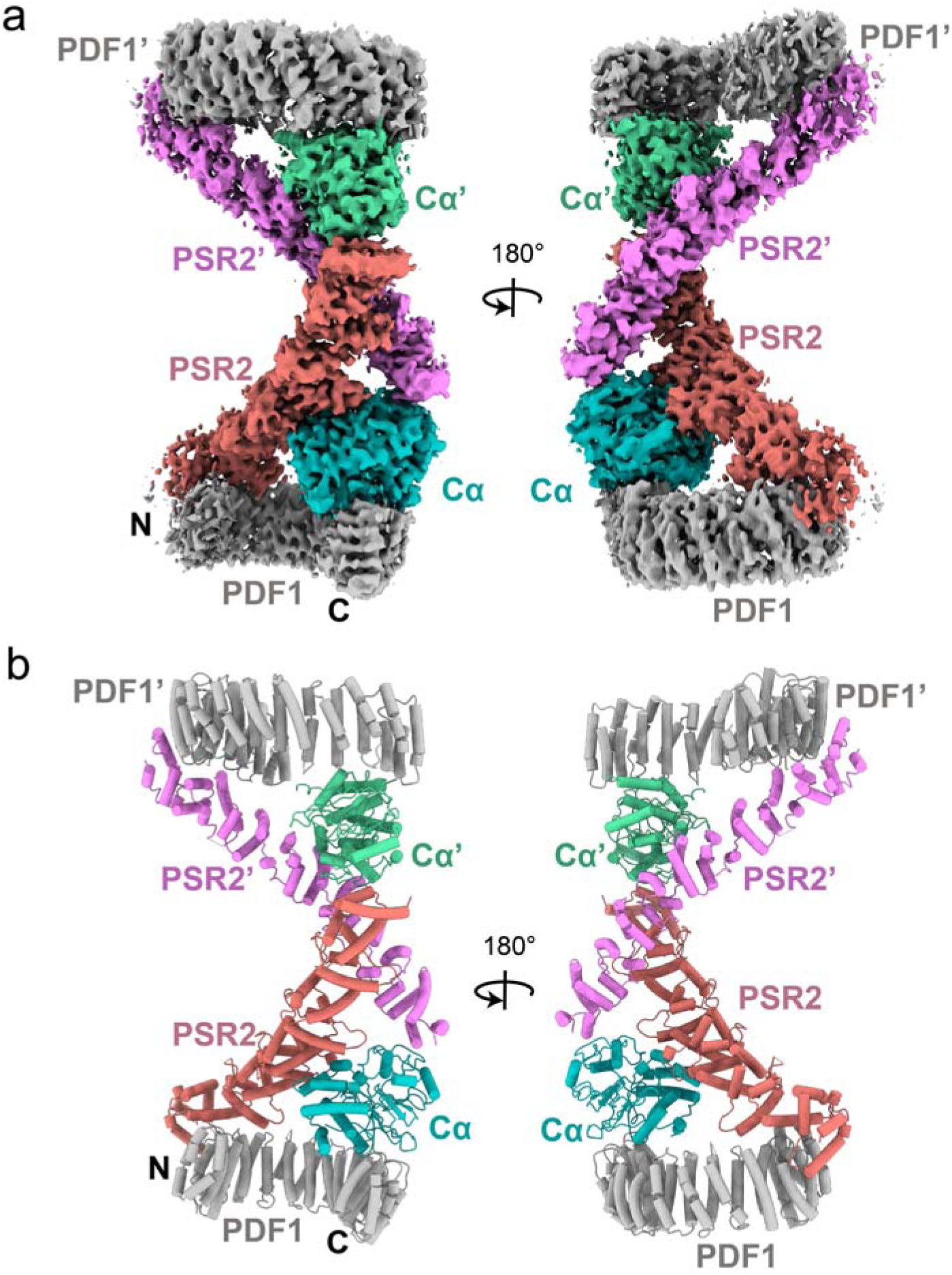
Overall cryo-EM structure of PSR2-PDF1-Cα hexamer complex. **a.** The final 3D reconstruction electronic density map of the PSR2-PDF1-Cα shown in two different orientations exists as a hexamer composing of two symmetrical heterotrimers (PSR2-PDF1-Cα and PSR2’-PDF1’-Cα’). N: the N-terminus of PDF1, which interacts with the N-terminus of PSR2; C the C-terminus of PDF1, which associates with Cα. **b.** The atomic model of the PSR2-PDF1-Cα shown in two different orientations exists as a hexamer composing of two symmetrical heterotrimers (PSR2-PDF1-Cα and PSR2’-PDF1’-Cα’).

The PSR2-PDF1-Cα hexamer is composed of two heterotrimers. Each PSR2 molecule binds to a PP2A core enzyme made up of one PDF1 and one Cα, forming a trimer. These trimers further dimerize to form a hexamer (Figs.1a and 1b). Unlike the known human PP2A holoenzymes, which are all single A-B-C heterotrimer (Supplementary Fig.3a)^14,15,30,31^. the PSR2-PDF1-Cα complex features two A-C core enzymes and two PSR2 effectors. The two rod-like PSR2 molecules intersect, forming a cross. Notably, The C-terminus of each PSR2 interacts with the Cα of the opposite heterotrimer, likely driving the dimerization.

### Interactions between PSR2 effector and the PP2A core enzyme

Within each heterotrimer, PDF1 adopts a horseshoe shape, similar to the published structures of human PP2A A subunit (PPP2R1A)^14,15,30,31^. PSR2 and Cα are positioned at the N- and C-termini of the PDF1 horseshoe, respectively (Fig.2a and Supplementary Fig.3b). This assembly resembles that of PP2A holoenzymes studied in human (Supplementary Fig.3a), wherein the C-terminal region of the A subunit interacts with Cα, forming a compact elliptical structure, and the N-terminal region interacts with B subunits. This similarity in structural configuration suggests that PSR2 functions as a molecular mimicry of PP2A B subunits and hijacks the PP2A core enzyme by substituting endogenous B subunits.

**Fig.2.**
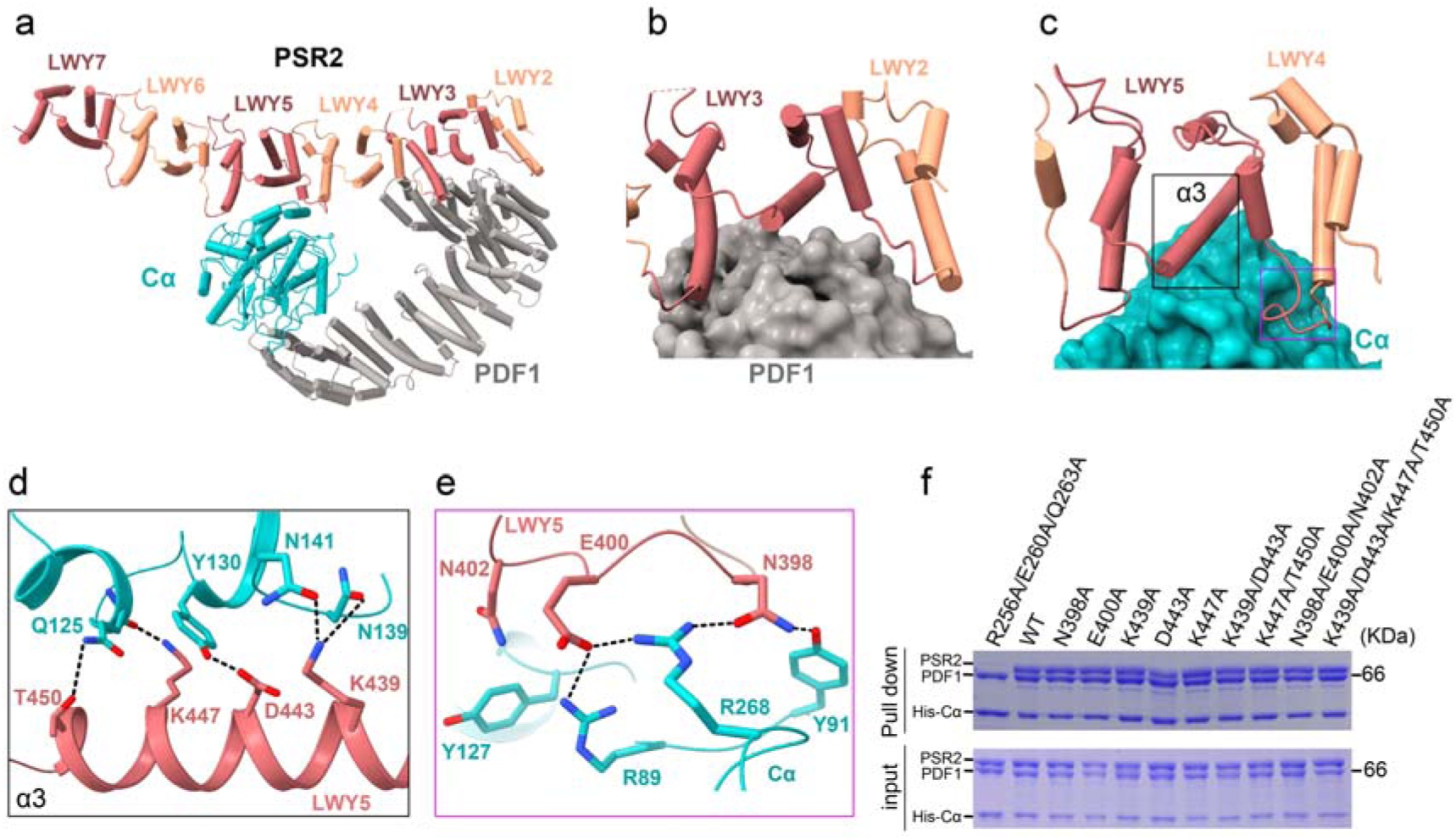
The interface formed by LWY4-LWY5 is not essential for PSR2 hijacking PP2A core enzyme. **a.** Overall structure of PSR2-PDF1-Cα heterotrimer holoenzyme separated from PSR2-PDF1-Cα hexamer holoenzyme. LWY modules are colored in two different colors (wheat and salmon) alternately. Because the electronic density of N-terminus of PSR2 is poor, we can’t resolve the atomic model of WY1 and partial LWY2. **b.** The groove-like structure formed by LWY2-LWY3 interacts with the N-terminus of PDF1. **c.** The groove-like structure formed by LWY4-LWY5 interacts with Cα. The third alpha helix of LWY3 is called α3. **d-e**. Seven residues in PSR2 are involved in interaction with Cα. **f**. The pull-down assays of PSR2 mutants in vitro show that the interface formed by LWY4-LWY5 is not essential for PSR2 to interact with PP2A core enzyme. The mutated location of PSR2^R256A/E260A/Q263A^ from LWY2-LWY3 (PP2A interacting module), our previous report indicate that this mutation almost abolishes its interaction with PDF1. Our current data show PSR2^R256A/E260A/Q263A^ also almost abolishes its interaction with PDF1-Cα.

The LWY_2_-LWY_3_ units of PSR2 anchor at the N-terminus of PDF1 (Fig.2b), similar to what was found in the crystal structure of PSR2-PDF1 complex using the N-terminal 390 aa fragment of PDF1^23^. In addition, in the PSR2-PDF1-Cα complex, we found that PSR2 also binds to Cα through the LWY_4_-LWY_5_ units (Figs.2a and 2c). The LWY4-LWY5 region of PSR2 is positioned on Cα, with the third α-helix of LWY_5_ (α3) forming the core interface (Fig.2c). In this interaction, the side chain of residue K439 forms hydrogen bonds with residues N139 and N141 in Cα (Fig.2d). Additionally, residues D443, K447, and T450 in α3 form hydrogen bonds with the Cα residues Y130 and Q125, respectively (Fig.2d). Furthermore, N398, E400, and N402 located between LWY4 and LWY5 further contribute to the interaction of PSR2 with Cα (Fig.2e).

The interaction of PSR2 and the PP2A catalytic subunit Cα is in contrast with the known human PP2A holoenzymes in which the B subunits had minimal or no interactions with Cα (Supplementary Fig.3a)^15,16,31,32^.We then evaluated the potential contribution of PSR2-Cα interaction to the assembly of the PSR2-PP2A complex. For this purpose, we mutated specific residues in LWY5 involved in the PSR2-Cα interaction in our structural model and assessed the mutants for PSR2-PDF1-Cα complex formation. Pull-down assays indicated that mutations of PSR2 potentially disrupting the interaction with Cα in the hexamer did not significantly hinder the complex assembly (Fig.2f). Furthermore, the quadruple mutant PSR2^K439A/D443A/K447A/T450A^ was co-precipitated with PP2A A subunits when expressed in *Nicotiana benthamiana*, and the protein complex exhibited phosphatase activity in a level that was comparable to that of wildtype PSR2 (Supplementary Fig.3c and 3d). These results suggest that the observed interactions of PSR2 with Cα through LWY4-LWY5 in the PSR2-PDF1-Cα complex is not essential for PSR2 to recruit PP2A core enzyme. This is consistent with the previous observation that PSR2-PP2A A interaction mediated by LWY2-LWY3 play a pivotal role in effector function (Fig.2f)^23^. Nonetheless, the interaction between PSR2 and Cα may facilitate the assembly and/or enhance the stability of the PSR2-PP2A complex during pathogen infection although this more subtle effect may not be detectable using purified or over-expressed proteins.

### LWY_7_-C**α** interactions drive PSR2-PP2A hexamer formation

In the PSR2-PDF1-Cα hexamer, the LWY_7_ unit at the C-terminus of PSR2 in one heterotrimer interacts with the Cα of the opposite trimer (hereafter Cα’), potentially facilitating the dimerization of two trimers to form a hexamer (Fig.3a). In this interaction interface, the fourth helix of LWY7 forms numerous hydrogen bonds and charged interactions with Cα’ (Figs.3b and 3c). Specifically, residues K636 and E647 of LWY7 form salt bridges with E192 and R206 of Cα’, while K646 and Q640 of LWY7 form hydrogen bonds with W209 and Y218 of Cα’ (Fig.3c).

**Fig.3.**
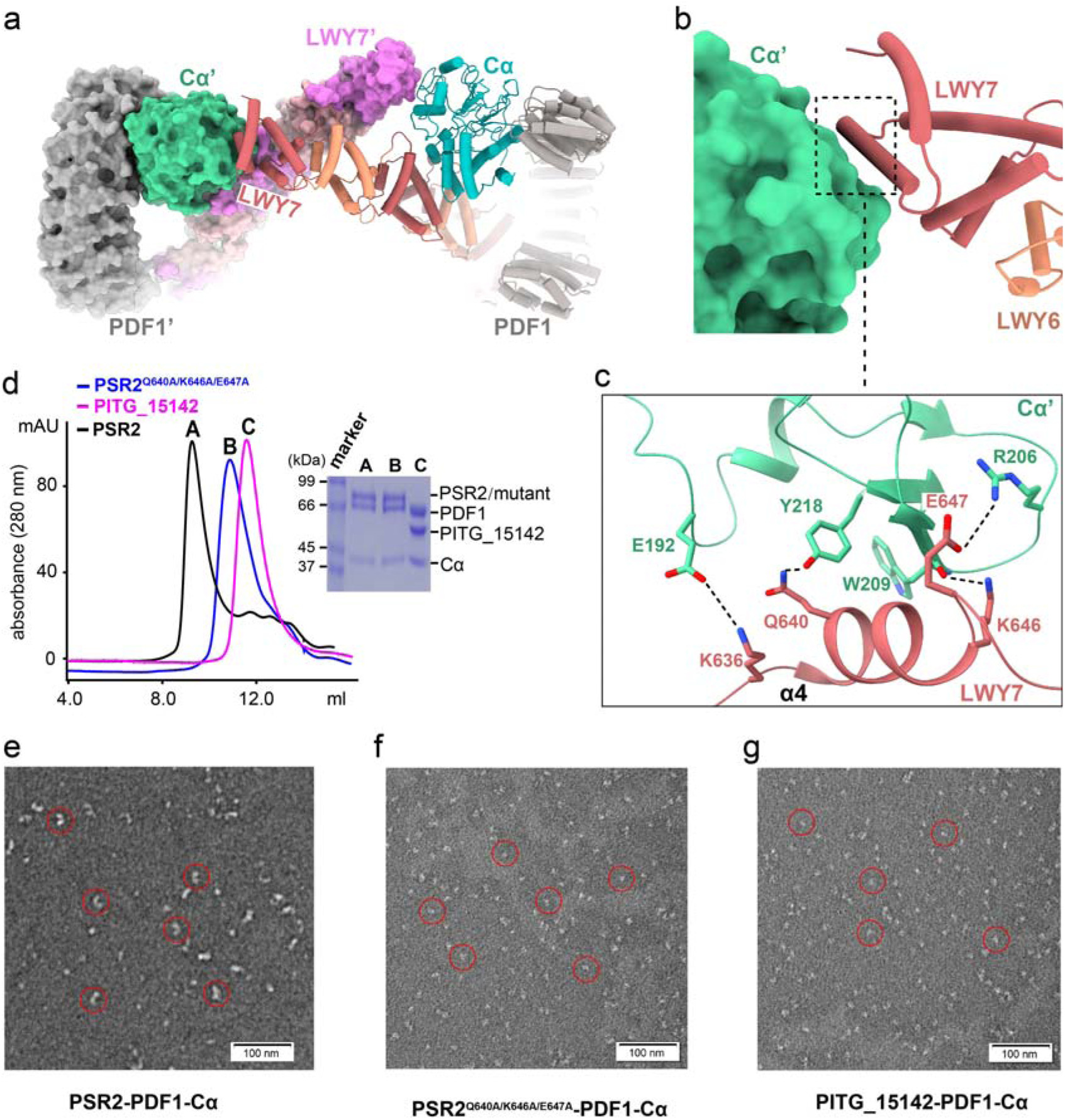
The interface formed by LWY7 module and C subunit is required for dimerization of two heterotrimers. **a.** Depiction of two symmetrical interfaces. LWY7 interacts with Cα’, and LWY7’ interacts with Cα. **b.** The fourth alpha helix of LWY7 contacts on the surface of Cα’. **c.** The interaction analysis between LWY7 and Cα’. There are four residues involved in interaction with Cα’. **d.** Gel filtration chromatography shows that PSR2^Q640A/K646A/E647A^ can still interact with PP2A core enzyme, but after binding to PP2A core enzyme the molecular weight shifts to the low level. In addition, the position of PITG_15142-PDF1-Cα ternary complex peak also shifts to the low level. **e-g**. Negative stain results of PSR2-PDF1-Cα (e), PSR2^Q640A/K646A/E647A^ -PDF1-Cα (f) and PITG_15142-PDF1-Cα (g) respectively, show that the particles of PSR2^Q640A/K646A/E647A^ -PDF1-Cα and PITG_15142-PDF1-Cα exist as heterotrimers.

To evaluate the significance of LWY7-Cα’ interaction in the formation of the PSR2-PP2A hexamer, we substituted Q640, K646 and E647 within the LWY_7_ with alanine. Gel filtration analyses revealed that PSR2^Q640A/K646A/E647A^ mutant formed a heterotrimeric complex with PDF1-Cα in vitro but failed to assemble into a hexamer (Fig.3d). Negative staining confirmed that the PSR2^Q640A/K646A/E647A^-PDF1-Cα complex particles were smaller than those form by PSR2-PDF1-Cα. These particles also exhibited a distinct shape, consistent with a disruption of heterotrimer dimerization by the mutations (Figs.3e and 3f), suggesting that LWY_7_ of PSR2 is essential for the hexamer formation.

In addition to PSR2, 12 *Phytophthora* effectors share a functional module similar to the LWY2-LWY3 of PSR2 also associate with plant PP2A core enzyme^23^. Among them, the *P. infestans* effector PITG_15142 has a WY1-(LWY)_4_ architecture and binds to PDF1 through its WY1-LWY2 units. However, PITG_15142 presumably lacks a C-terminal LWY unit equivalent to LWY7 of PSR2. We therefore examined whether PITG_15142 could form a hexamer with the PP2A core enzyme. Gel filtration and negative staining showed that PITG_15142 only formed a heterotrimeric complex with PDF1-Cα, with a size and shape similar to the PSR2^Q640A/K646A/E647A^-PDF1-Cα complex (Figs.3d and 3g). These results demonstrate that LWY7 of PSR2 is essential for the PSR2-PP2A hexamer formation through mediating the dimerization of two heterotrimers.

### Structural dynamics of trimeric PSR2-PP2A complexes

To investigate the functional advantages of the hexameric PSR2-PP2A holoenzymes compared to a canonical heterotrimer, we reconstituted the complex using the PSR2 mutants PSR2^Q640A/K646A/E647A^ and PSR2^ΔWY1+ΔLWY^^7^, the latter lacking both LWY7 and LWY1 units (Supplementary Fig.4). Among the cryo-EM samples tested, only the PSR2^ΔWY1+ΔLWY7^-PDF1-Cα was suitable for data collection. Two distinct cryo-EM structures of the trimeric form (designated Trimer-S1 and Trimer-S2) were determined with resolutions of 4.5 Å and 4.4 Å, respectively (Figs.4a, 4b, Supplementary Fig.5 and Table 1).

**Fig.4.**
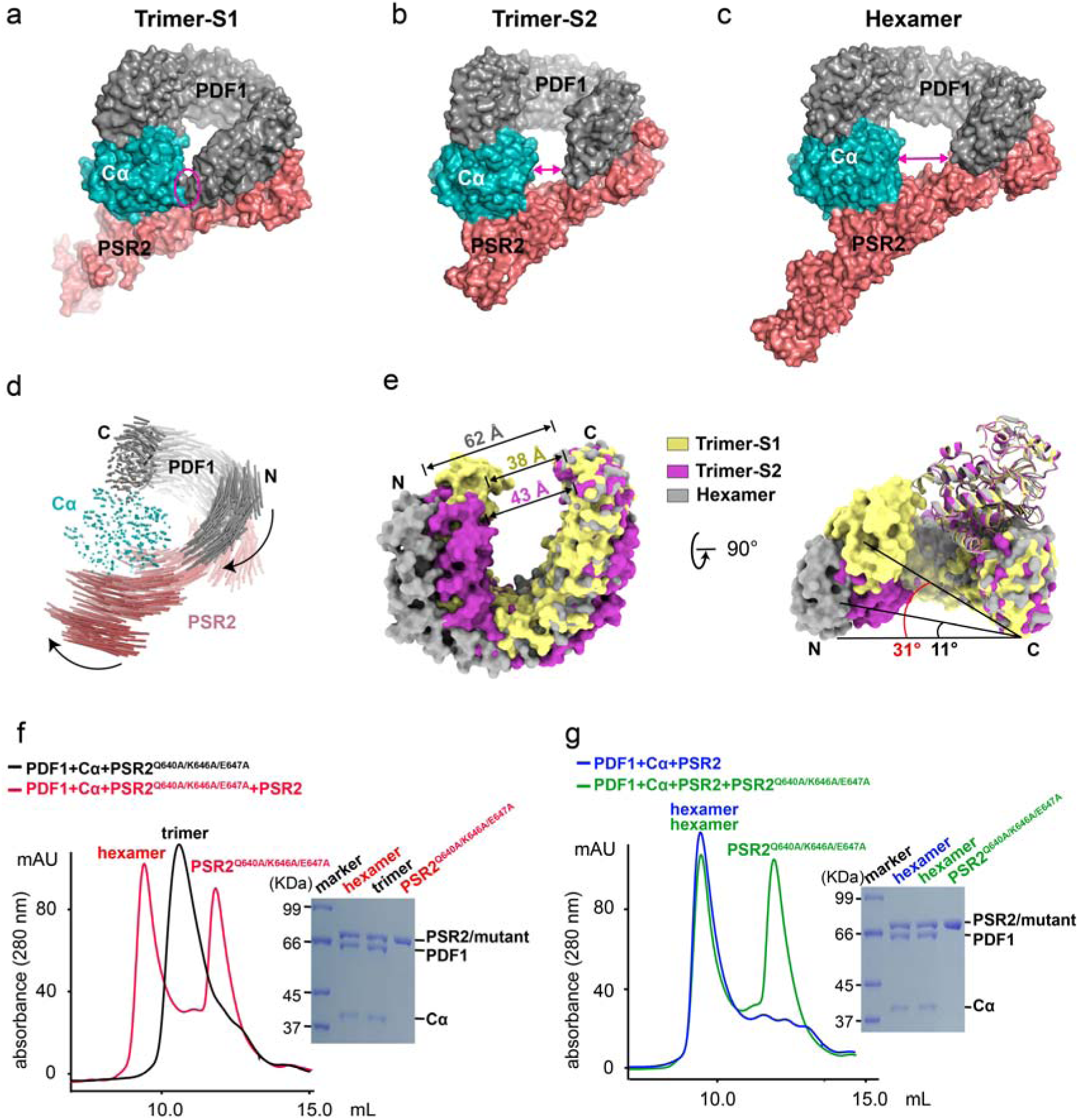
PSR2-PDF1-Cα complex in heterotrimeric state possesses two main conformations. **a.** Surface representation view of overall structure of PSR2-PDF1-Cα heterotrimer holoenzyme in Trimer-S1. **b.** Surface representation view of overall structure of PSR2-PDF1-Cα heterotrimer holoenzyme in Trimer-S2. **c.** Surface representation view of overall structure of PSR2-PDF1-Cα heterotrimer holoenzyme separated from PSR2-PDF1-Cα hexamer holoenzyme. **d.** Structural superposition shows the structural transition from trimer-S2 to trimer-S1. Vector length correlates with the transition scale. N: the N-terminus of PDF1, which interacts with the N-terminus of PSR2; C: the C-terminus of PDF1, which associates with Cα. **e.** Structural comparison of PDF1 from hexamer, Trimer-S1 and Trimer-S2 shows that the conformations of PDF1 in different states are different. N: the N-terminus of PDF1, which interacts with the N-terminus of PSR2; C: the C-terminus of PDF1, which associates with Cα. **f.** PSR2 efficiently displaced PPSR2^Q640A/K646A/E647A^ from a preformed PPSR2^Q640A/K646A/E647A^ -PDF1-Cα heterotrimeric holoenzyme in vitro. Gel filtration chromatography shows two peaks representing the PSR2-PDF1-Cα hexameric holoenzyme and excess PSR2^Q640A/K646A/E647A^ dropped from the preformed PPSR2^Q640A/K646A/E647A^ -PDF1-Cα holoenzyme. Gel filtration chromatography of the PPSR2^Q640A/K646A/E647A^ -PDF1-Cα heterotrimeric holoenzyme was used as a control. **g.** PSR2^Q640A/K646A/E647A^ could not efficiently displace PSR2 in a preformed PDF1-PSR2-Cα hexameric holoenzyme. Gel filtration chromatography of the PDF1-PSR2-Cα hexameric holoenzyme was used as a control.

In both trimeric conformations, one PSR2^ΔWY1+ΔLWY7^ binds to one PDF1 and one Cα, forming a trimer with PSR2 and Cα positioned at the N- and C- termini of PDF1 (Figs 4a, 4b and Videos S1–S2), respectively. This arrangement resembles configuration of the PSR2-PDF1-Cα hexamer (Fig.4c). However, superposition of the Cα proteins in the two trimeric states revealed that PDF1 and PSR2^ΔWY1+ΔLWY7^ adopt different conformations in Trimer-S1 and Trimer-S2 (Fig.4d). Notably, the Cα almost connects the N- and C-termini of PDF1 in Trimer-S1 (Figs.4a and 4b).

PDF1 displays a partially elliptical structure, with varying size and shape in the two trimeric and one hexameric states (Fig.4e). The distances between the N- and C-termini of PDF1 are 38 Å in Trimer-S1, 43 Å in Trimer-S2, and 62 Å in the hexamer. In the trimeric states, PDF1 adopts a narrower, more elongated elliptical conformation compared to the hexamer. Additionally, the N-terminal region of PDF1 exhibits significant tilting, with angles of 31° and 11° in Trimer-S1 and Trimer-S2, respectively (Fig.4e), and Cα nearly bridges the N- and C-termini of PDF1 in Trimer-S1.

The conformational variability and weaker electron density in these trimeric states suggest that the N-terminal region of PDF1 is highly flexible and dynamic. These observations underscore the structural adaptability of the trimeric complexes, contrasting with the more rigid architecture of the hexamer.

### Hexametric assembly increases the stability of PSR2-PP2A holoenzyme

The dynamics of PDF1 influence the movement of the PSR2 molecule bound to its N-terminus (Video S3), leading to conformational variability in the trimeric PSR2-PP2A complexes. In contrast, in the hexameric form, interactions between LWY7 of PSR2 and Cα’ constrain the C-terminal movements of PSR2, enhancing the stability of the holoenzyme (Fig.3a).

To compare the stability of the hexameric and trimeric complexes, we performed a competition assay. When wild-type PSR2 was introduced into a pre-formed PSR2^Q640A/K646A/E647A^-PDF1-Cα complex, it displaced PSR2^Q640A/K646A/E647A^ mutant, forming hexameric complex with PDF1-Cα (Fig.4f). In contrast, adding PSR2^Q640A/K646A/E647A^ mutant to a pre-formed PSR2-PDF1-Cα complex did not induce an efficient switch to the trimeric state (Fig.4g). These results further confirm that the hexameric complex is more stable than its trimeric counterpart, and the interaction between the two trimers likely stabilizes the overall hexameric assembly (Fig.3a).

### Dimerization of PSR2-PP2A heterotrimers opens the catalytic pocket

To evaluate the functional implications of hexamerization, we compared the trimeric and hexameric structures, focusing on the accessibility of the catalytic pocket in Cα. In the trimeric states, the C-terminal region of PSR2 exhibits significant flexibility, with the LWY4-LWY5 region dynamically sliding toward Cα (Video S3). This movement causes the catalytic pocket to be fully covered in Trimer-S1 and partially covered in Trimer-S2 (Figs.5a and 5b).

**Fig.5.**
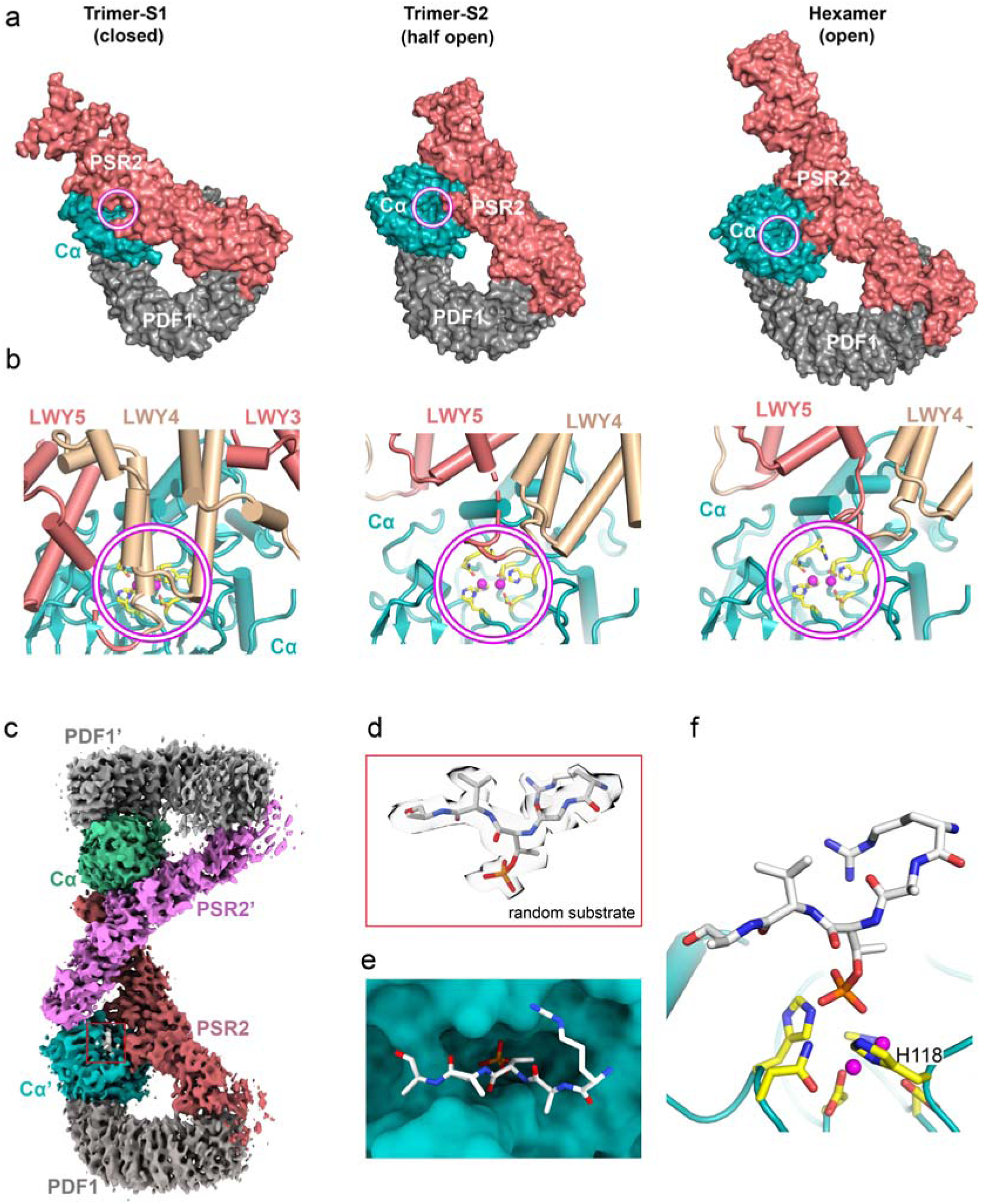
The catalytic pocket of Cα in hexameric state is more open. **a.** Surface representation view of overall structure of PSR2-PDF1-Cα holoenzyme in Trimer-S1, Trimer-S2 and Hexamer show that the relative positions between PSR2 and the catalytic pocket of Cα. The red circles are the catalytic pockets of Cα. **b.** Detailed cartoon representation of the relative positions between Cα and LWY4-LWY5 in Trimer-S1, Trimer-S2 and Hexamer. **c.** Overall structure of PSR2^hexamer^-PDF1-Cα^substrate^ hexamer holoenzyme binding to random substrates. The region marked by red rectangle is the catalytic pocket of C subunit binding to the random substrate. **d.** Electronic density of random substrate from PSR2^hexamer^-PDF1-Cα^substrate^ hexamer holoenzyme map. **e.** Surface representation views of random substrate binding to catalytic pocket of C subunit. **f.** Cartoon representation views of random substrate binding to catalytic pocket of C subunit.

In contrast, in the hexameric state, interactions between LWY7 of PSR2 and Cα’ from the opposite trimer stabilize PSR2 and keep it away from the catalytic pocket, leaving the pocket completely open (Figs.5a and 5b). These results suggest that in the trimeric states, PSR2 may restrict substrate access to the catalytic pocket, while dimerization in the hexamer enhances catalytic pocket accessibility via opening the catalytic pockets, potentially improving the efficiency of substrate entry.

### Substrate bound in the catalytic pocket of C**α** subunit

To further confirm that the PSR2-PDF1-Cα hexamer has open catalytic pockets, we incubated a synthetic phosphopeptide (RRA(pT)VA) with the PSR2-PDF1-Cα complex as a substrate. We determined cryo-EM structure of this PP2A holoenzyme-substrate complex at a resolution of 3.39 Å (Fig.5c and Supplementary Fig.6). The complex retained the hexameric arrangement, with the catalytic pockets of Cα and Cα’ containing clear extra electron density, enabling the unambiguous docking of one phosphopeptide molecule into each pocket (Figs.5c and 5d). The phosphopeptide is positioned in the cleft of Cα (Fig.5e), with its phosphate group being located adjacent to the catalytic residue H118 (Fig.5f)^32^, mimicking the substrate-binding state. Interestingly, the binding position of the phosphopeptide mirrors that of microcystin-LR, a competitive inhibitor of PP2A complex (Supplementary Fig.7a)^13,33,34^. This similarity validates the substrate-bound state of the PSR2-PP2A hexamer.

The structure of the substrate-bound hexamer displayed a similar arrangement to the unbound form (Supplementary Fig.7b). In this structure, the LWY4-LWY5 region of PSR2 remained distant from the Cα catalytic pocket, maintaining the open conformation of the catalytic pockets, as seen in the substrate-free hexamer (Fig.5a and Supplementary Fig.7c). In contrast, in the trimeric forms, the LWY4-LWY5 region of PSR2 either completely or partially covered the Cα catalytic pocket (Supplementary Figs.7d and 7e). These findings suggest that hexamerization of the PSR2-PP2A complex promotes the opening of the catalytic pocket, whereas the trimeric state may hinder substrate entry into the catalytic pocket.

### Hexamerization of PSR2-PP2A holoenzyme is important for PSR2 virulence activity

To examine the functional significance of the hexameric form of the PSR2-PP2A holoenzyme, we generated transgenic *A. thaliana* plants expressing either PSR2 or PSR2^Q640A/K646A/E647A^. For each construct, multiple independent transgenic lines with comparable protein expression levels were evaluated for susceptibility to *Phytophthora capsici* (Supplementary Fig.8). *A. thaliana* plants expressing wild-type PSR2 showed increased susceptibility to *P. capsici* (Figs.6a and 6b), consistent with our previous studies^23,35^. Although expression of PSR2^Q640A/K646A/E647A^ in *A. thaliana* also led to hypersusceptibility compared to wildtype plants, these transgenic plants were significantly less susceptible than those expressing wild-type PSR2 (Figs.6a and 6b). These results indicate that PSR2 has a higher virulence activity when forms the hexameric holoenzyme with the PP2A core enzyme than the trimeric form.

**Fig.6.**
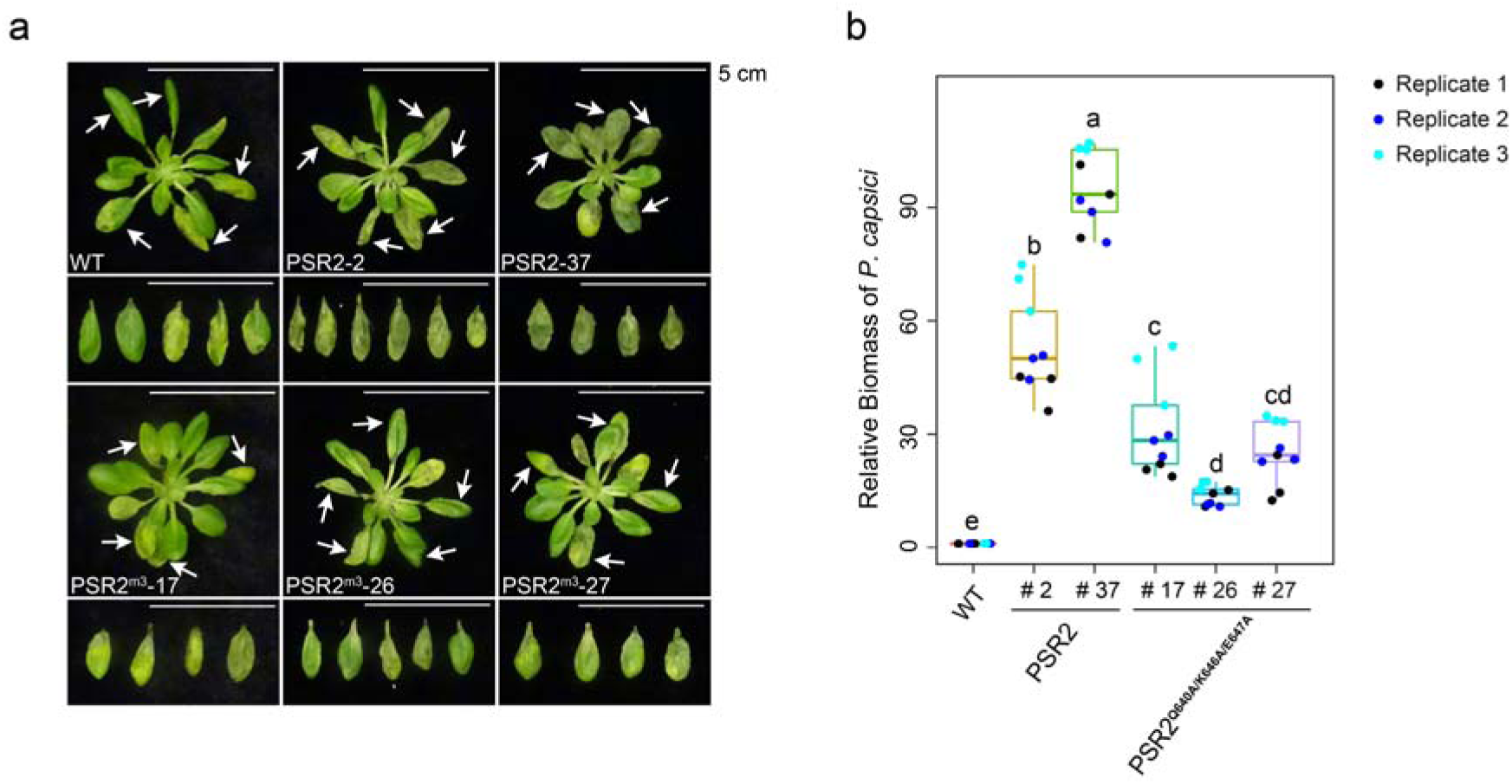
The hexameric assembly of PSR2 with plant PP2A core enzyme is important for its virulence function. **a.** The virulence activity of PSR2 is higher than PSR2^Q640A/K646A/E647A^ (PSR2^m3^). Four-week-old Arabidopsis plants were inoculated with *Phytophthora capsici* isolate LT263. Photos were taken at 2 days post-inoculation (dpi) with zoospore suspensions. Arrows indicate inoculated leaves. WT, wild-type Col-0. **b.** Relative biomass of *P. capsici* in inoculated plants at 2 dpi was determined by qPCR (n ≥ 20 in each sample per experiment). Data from three independent replicates are presented. One-way ANOVA and post hoc Tukey were used for statistical analysis. Different letters label significant differences (p < 0.05).

## Discussion

Pathogen effectors are key virulence proteins that directly manipulate host cellular processes to promote diseases. The most common host targets are proteins, and the protein-protein interactions with specific host target are critical for the virulence activity of the effectors. However, molecular details of the structure-function relationship of effector-target complexes are rarely understood.

In this study, we described an effector-target complex, which represents a unique structure of PP2A holoenzyme. The protein phosphatase PP2A functions as a holoenzyme with three different subunits. The core enzyme is highly conserved across the biological kingdoms. Importantly, PP2A core enzyme is commonly targeted by oncogenic viruses in animals, bacterial and *Phytophthora* pathogens in plants, demonstrating its importance in pathogenesis^16–23^. In particular, PP2A core enzyme can be hijacked by *Phytophthora* effectors, potentially serve as functional mimics of the substrate-binding subunit in the holoenzyme^23^. Here, we solved in the Cryo-EM structure of one of such *Phytophthora* effectors, PSR2, in complex with PP2A core enzyme consisting of a plant A subunit and a human C subunit. This complex forms a novel structure that differs from the known PP2A holoenzymes. Although PSR2 interacts with the A and C subunits of the PP2A core enzyme similarly to endogenous B subunits, its interaction with the C subunit of the opposite core enzyme through a C-terminal region of PSR2 enables the formation of a hexameric holoenzyme. This hexameric assembly not only increases the stability of the PSR2-PP2A complex but also opens the catalytic pocket within the C subunit, potentially facilitating substrate entry and enhancing the virulence function of PSR2. This is consistent with a role of PSR2 that can efficiently hijack the core enzyme from the host.

PSR2 and other PP2A core enzyme-interacting *Phytophthora* effectors are composed of (L)WY tandem repeats^23^. The (L)WY units are structurally conserved but functionally variable^27^. Distinct (L)WY unit or unit combinations can mediate specific interactions with host targets, thus serving as functional modules. Combining different modules is a promising mechanism to promote functional diversification in an effector repertoire^36,37^. The effectors that hijack the host PP2A core enzyme share the same N-terminal (L)WY-LWY module that mediates interaction with the PP2A A subunit but their C-terminal (L)WY units are variable. We show that the C-terminal LWY7 of PSR2 is critical for forming the hexameric assembly of PSR2-PP2A holoenzyme. A similar LWY unit is absent from most of the other PP2A-interacting effectors. An interesting hypothesis is that the LWY7 was introduced to PSR2 through shuffling of (L)WY unit(s), enhancing its association with the host PP2A core enzyme and thus virulence activity.

In summary, our integrative structural and functional analysis of a pathogen effector in a complex with its host target advances a fundamental understanding of the molecular mechanisms underlying microbial pathogenesis. The knowledge of effector-PP2A holoenzymes has broad implications given the important role of PP2A as a hub that is commonly targeted by various pathogens of human and plants. The molecular details at the effector-PP2A interaction interface offers opportunities to develop novel management strategies of the devastating *Phytophthora* diseases.

### Data availability

The atomic coordinates of the PSR2^hexamer^-PDF1-Cα, PSR2^hexamer^-PDF1-Cα^substrate^, Trimer-S1 and Trimer-S2 have been deposited in the Protein Data Bank, and the accession codes are 9J5I, 9J5K, 9J5R and 9J5N respectively. The overall cryo-EM map of PSR2^hexamer^-PDF1-Cα, PSR2^hexamer^-PDF1-Cα^substrate^ Trimer-S1 and Trimer-S2 have been deposited in the Electron Microscopy Data Bank, and the accession codes are EMD-61142, EMD-61144, EMD-61150 and EMD-61147, respectively.

## Acknowledgements

We thank staff of the Center for Biological Imaging, Core Facilities for Protein Science at the Institute of Biophysics (IBP), Chinese Academy of Sciences (CAS). We thank Cong Yu for kindly providing Hi-5 insect cell, Long Si for assistance of cryo-EM sample preparation, and Yao Rong for help of insect cell culture. We also thank the tissue culture support team at the Sainsbury Laboratory for generating the transgenic plants. **Funding:** Y.W. is supported by grants from Beijing Municipal Science & Technology Commission (Z231100007223004), Beijing Natural Science Foundation (5232022), the Natural Science Foundation of China (32330055, 31930065, 22121003, 32071198, 32071444), National Key R&D Program of China (2023YFC3403400, 2023YFA0915000), the Chinese Academy of Sciences (XDB0570000 and XDB37010202). W. M. is supported by Gatsby Charitable Foundation, UKRI BBSRC Grants BB/W016788/1 and BBS/E/J/000PR9797.

## Author contributions

Y.W. and W.M conceived the project, guided the execution of the experiments, and oversaw the project. Jinlong Wang performed the proteins purification and solved the all structures. Jinlong Wang, H.L., W.S., Jiuyu Wang, X. F., X.Y., C.L. and G.S. did the experiments and analyzed the data. Jinlong Wang and H.L. prepared figures and table. Y.W., W.M., Jinlong Wang and H.L. wrote the manuscript with contributions from all authors.

## Declaration of interests

The authors declare no competing interests.

## Method details

### Plant materials and growth conditions

*Arabidopsis thaliana* used for generating transgenic plants and *Nicotiana benthamiana* plants were grown in a growth room at 22°C with a 16/8h light/dark regime. Arabidopsis used for *Phytophthora capsici* infection assay were grown in a growth room at 22°C with an 8/16h light/dark regime. Sterile Arabidopsis seedlings were grown on plates containing Murashige-Skoog medium and 1% sucrose supplemented with 0.8% Phytagel in a growth chamber with the setting of 22°C and a 16/8 h light/dark regime.

#### Phytophthora capsici

*P. capsici* isolate LT263 was used in this study and was grown on fresh 10% V8 plates at 25°C in the dark for mycelia growth. Zoospores release was based on a method descried in Method Details.

### Plasmid construction

PSR2 or mutated PSR2 was cloned without their N-terminal secretion signal peptide for various experiments. For protein expression or generating stable transgenic plants, PSR2 were cloned into the pENTR/D-TOPO vector (Invitrogen) and then destination vectors (pEarleyGate100 or pEarleyGate101, Invitrogen) using the LR Clonase II-based gateway cloning system.

### Generation of Arabidopsis transgenic plants

To obtain the transgenic plants, *Agrobacterium tumefaciens* carrying the corresponding construct was used for Arabidopsis transformation. The resulting transgenic T1 seeds were screened on ½MS medium with Phosphinothricin (Duchefa) or germination medium with Kanamycin. The PSR2 or its mutant variant was detected using anti-PSR2 antibody through western blotting^35^.

### Immunoprecipitation of PSR2 or its mutant variant complexes

Interaction of PSR2 or PSR2^K439A/D443A/K447A/T450A^ with PP2A A subunits were also detected in *N. benthamiana*. A. tumefaciens carrying constructs for expressing PSR2 or its mutant variant were infiltrated into leaves of 3-week-old plants. One grams of infiltrated tissues were ground in liquid nitrogen and suspended in 1 mL IP buffer (10% (v/v) Glycerol, 50 mM Tris-HCl pH 7.5, 50 mM NaCl, 1 mM EDTA, 1×protease inhibitor mixture, 5 mM DTT, 2% PVPP and 0.1% NP-40). Samples were centrifuged at 13000 rpm for 15 min at 4°C. Supernatants were incubated with 10 μL of anti-Flag magnetic beads and incubated for 2 hr at 4°C. Beads were washed three times (10% (v/v) Glycerol, 50 mM Tris-HCl pH 7.5, 50 mM NaCl, 1 mM EDTA, and 0.1% NP-40). Bound PSR2 or its mutant variant and their associated proteins were separated by SDS-PAGE electrophoresis for western blotting. PSR2 or PP2A A subunits were detected using anti-Flag and anti-PP2A A antibodies respectively.

### PP2A phosphatase activity assay

To measure the phosphatase activity of PSR2 or PSR2^K439A/D443A/K447A/T450A^ complexes in *N. benthamiana*, they were transiently expressed in leaves using Agro-infiltration and enriched from leaf tissues using anti-Flag magnetic beads as described above. PP2A phosphatase activity was measured using a non-radioactive molybdate dye-based phosphatase assay kit (#V2460, Promega) in which a synthetic phosphopeptide (RRA[pT]VA) was used as the substrate. A reaction mixture containing 100 μM phosphopeptide and immunoprecipitated proteins was incubated at 37°C for 5 min with or without 1 nM Okadaic acid, which is a PP2A-specific inhibitor^38^. An equal volume of molybdate dye-additive was used to stop the reaction. Phosphate released from the phosphopeptide was measured as absorbance at 600 nm against a standard curve. Relative PP2A activity was calculated as the ratio between the experimental group and the control (the leaves expressing infiltrated with the empty vector).

### Arabidopsis infection assays by *Phytophthora capsici*

Inoculation using *P. capsici* isolate LT263 in Arabidopsis was performed as previously described^39^. LT263 was grown on 10% V8 medium at 25°C in the dark until mycelia covered the whole plate. To induce sporulation, mycelium plugs were first washed and then incubated with sterile tap water at 25°C for 24 hours in the dark. Zoospore release was induced by 4°C incubation for 40 min, followed by light induction for 20 min at room temperature. Zoospores were collected using one layer of miracloth (Millipore) for making suspensions (200–500 zoospores/μL) to be used for inoculation. 3-6 adult rosette leaves per Arabidopsis plant were inoculated using ∼20 μL zoospore suspension applied to the abaxial side of each leaf. Leaves treated with water were used as a mock control. The inoculated plants were placed in a growth chamber with a transparent cover to keep high humidity. Disease symptoms were monitored three days after inoculation and DNA was extracted from all the inoculated leaves (n ≥ 20 per treatment). The biomass of *P. capsici* was determined by qPCR using *P. capsici* specific primers. The Arabidopsis *rub4* was used an internal control. Relative biomass of *P. capsici* in mutant or transgenic plants was determined by comparing to the value from the biomass in wildtype (WT) plants, which was set as “1”.

### Protein expression and purification

PSR2 (encoding residues 18-670) was cloned into an expression vector pRSFDuet-sumo with a 6 His-Sumo tag at the N terminus. PDF1 (encoding residues 1-587) were cloned into an expression vector pRSFDuet with a 6 His tag or 6 His-Sumo tag at the N terminus. PITG_15142 (encoding residues 23-490) were cloned into an expression vector pET28a-sumo with a 6 His sumo tag at the N terminus. All proteins, mutants and truncations were overexpressed in *E. coli* BL21 (DE3) (Novagen) cells and were induced by adding 0.1 mM IPTG with the OD_600_ = 0.4-0.8 at 16 °C for 16 hr. Cells were lysed by sonication in buffer containing 20 mM Tris-HCl (pH 7.5) and 0.5 M NaCl at 4°C, and target proteins were purified by Ni-NTA Sepharose resin (GE Healthcare), and further fractionated by ion exchange (GE Healthcare). Proteins with His-Sumo tag needed ubiquitin-like protein 1 (Ulp1) protease to cleave their His-Sumo tag and dialyzed against the buffer containing 20 mM Tris-HCl (pH 7.5), 0.3 M NaCl for 2 hr at 4 °C, afterward, the proteins were further purified by Ni-NTA Sepharose resin again to remove excised His-Sumo tag and then fractionated by ion exchange. Cα with 8 His tag was overexpressed in baculovirus-infected Hi-5 suspension culture, purified by Ni-NTA Sepharose resin (GE Healthcare), and further fractionated by ion exchange (GE Healthcare). Detailed protein purification methods of Cα were described previously^15^.

### Reconstitution of PSR2-PDF1-C**α** and PITG_15142-PDF1-C**α** complexes

To prepare the PSR2-PDF1-Cα and PITG_15142-PDF1-Cα complexes, purified PDF1, Cα and PSR2/PSR2 mutations/PSR2 truncations/PITG_15142 were mixed in the buffer containing 20 mM Tris-HCl pH 7.5, 1mM DTT, 300 mM NaCl at a molar ratio of 1: 1: 1. Components were added in the order listed above and incubated for 15 min on ice before adding the next component. The resulting complex was purified by gel filtration chromatography (Superdex 200 increase 10/300GL, GE Healthcare) and analyzed by using SDS-PAGE.

### Pull-down assay

The purified 8 His tag-fusion Cα was mixed with untagged PDF1 at a molar ratio of 1: 1 in buffer (20 mM Tris-HCl pH 7.5,100 mM NaCl, 3 mM β-ME) on ice for 15 min, then adding equal molar PSR2 and incubating for 15 min on ice too. Protein mixture was added to Ni-NTA Sepharose resin (GE Healthcare). The non-specifically bound proteins were removed by excess wash buffer (20 mM Tris-HCl pH 7.5, 100 mM NaCl, 40 mM imidazole). The bound proteins were eluted by elution buffer (20 mM Tris-HCl pH 7.5, 100 mM NaCl, 500 mM imidazole). The flow-through collections were analyzed by SDS-PAGE and stained by Coomassie Brilliant Blue.

### Competition experiments

A Superdex 200 Increase 10/300 gel filtration column (GE Healthcare) was used to analyze the state of protein complexes at a flow rate of 0.2 mL/min and with the absorbance monitored at 280 nm. Preformed PSR2^Q640A/K646A/E647A^-PDF1-Cα complexes were prepared through incubating purified PDF1, Cα and PSR2^Q640A/K646A/E647A^ at a molar ratio of 1: 1: 1. Components were added in the order listed and incubated for 15 min on ice before adding the next component. Then, equimolar PSR2 proteins were added to sample and incubated for another 15 min on ice. 100 μL of the sample was applied to pre-equilibrated column (20 mM Tris-HCl pH 7.5, 300 mM NaCl, 1 mM DTT) and analyzed by gel filtration chromatography. Fractions were also analyzed by 12% SDS-PAGE gel and visualized by Coomassie blue staining. We also tested whether PSR2^Q640A/K646A/E647A^ could facilitate the dissociation of PSR2 from a preformed PSR2-PDF1-Cα complex using this similar procedure.

### Cryo-EM Specimens Preparation and data collection

3.5 μL droplets of PSR2-PDF1-Cα and PSR2^ΔWY1+ΔLWY7^-PDF1-Cα complex samples were applied onto glow-discharged 1.2/1.3 Au 300 mesh grid (Ni/Ti; Lehua Electronic Technology) at a concentration of 0.3 mg/ml, respectively. After incubation for 10 s, excess sample was blotted with filter paper for 2 s-5 s in 100% humidity at 4°C and the grid was plunged into liquid ethane by using Vitrobot Mark IV (Thermo Fisher Scientific). The grids samples were named PSR2^hexamer^-PDF1-Cα and PSR2^heterotrimer^-PDF1-Cα, respectively and were stored in liquid nitrogen before being checked by the electron microscopy equipment.

In order to capture the state that PSR2^hexamer^-PDF1-Cα complex binds to the substrate, we use a synthetic phosphopeptide (RRA[pT]VA) as a random substrate. A final concentration of 1 mM synthetic phosphopeptide was added to the 0.3 mg/ml PSR2^hexamer^-PDF1-Cα complex and then 3.5 μL droplets of PSR2^hexamer^-PDF1-Cα complex incubating with synthetic phosphopeptide was applied onto glow-discharged 1.2/1.3 Au 300 mesh grid. After incubation for 5 s, excess sample was blotted with filter paper for 2 s-5 s in 100% humidity at 4°C and the grid was plunged into liquid ethane by using Vitrobot Mark IV. The grids sample was named PSR2^hexamer^-PDF1-Cα^substrate^ was stored in liquid nitrogen before being checked by the electron microscopy equipment.

The cryo-EM data of different samples were collected by using 300 kV FEI Titan Krios electron microscope equipped with Gatan K2 Summit direct electron detector positioned post a GIF quantum energy filter with energy filtered mode, the camera was in super-resolution mode and the physical pixel size is 1.04 Å (0.52 Å super-resolution pixel size). The defocus ranges from 1.2 to 1.8 μm and each image was exposed for 8 s, resulting in a total dose of 50 e^-1^/Å^2^ and 32 frames per movie stack. Automatic data collection were facilitated by using Serial EM software^40,41^.

### Image Processing and model building

The movies stack of different cryo-EM samples (PSR2^hexamer^-PDF1-Cα/PSR2^heterotrimer^-PDF1-Cα/PSR2^hexamer^-PDF1-Cα^substrate^) were imported into respective project files of CryoSPARC and then were processed one by one by using patch motion correction and patch CTF (contrast transfer function) estimation^42^. Micrographs with low quality were excluded during executing manual curation. Particles were manually picked and processed with reference-free 2D classification. 2D class average images were selected as references for automatic particle picking of the entire dataset. The total particles were picked and processed by reference 2D classification. Choosing particles with good quality for further Homogeneous Refinement and Non-uniform Refinement^42^. The detailed procedures were illustrated in flowchart (Supplementary Figs.2, 5 and 6). All structures were built manually in COOT^43^, and structural refinement were performed using PHENIX^44^. All structural figures were prepared using PyMOL (https://www.pymol.org/) or UCSF ChimeraX^45^.

### Quantification and statistical analysis

Data are represented as the mean ± s.e.m. or as box-and-whisker plots in which the center line indicates the median, the bounds of the box indicate the upper and lower quantiles, using R Studio (https://www.r-project.org/) or GraphPad Prism 8.0. Statistical analyses performed using GraphPad Prism 8.0. Relative PP2A activities were analyzed using a two-tailed Student t-test. Data for testing the significant differences between Arabidopsis genotypes were performed using One-way ANOVA and post hoc Tukey.

## Notes

### Competing Interest Statement

The authors have declared no competing interest.

## References

1 Jones, J. D. G. & Dangl, J. L. The plant immune system. Nature 444, 323–329 (2006). 10.1038/nature05286

2 Wang, Y., Pruitt, R. N., Nürnberger, T. & Wang, Y. Evasion of plant immunity by microbial pathogens. Nature Reviews Microbiology 20, 449–464 (2022). 10.1038/s41579-022-00710-3

3 Jones, J. D. G., Staskawicz, B. J. & Dangl, J. L. The plant immune system: From discovery to deployment. Cell 187, 2095–2116 (2024). 10.1016/j.cell.2024.03.045

4 Nürnberger, T., Brunner, F., Kemmerling, B. & Piater, L. Innate immunity in plants and animals: striking similarities and obvious differences. Immunological Reviews 198, 249–266 (2004). 10.1111/j.0105-2896.2004.0119.x

5 Brautigan, D. L. Protein Ser/ϑThr phosphatases – the ugly ducklings of cell signalling. The FEBS journal 280, 324–325 (2013). 10.1111/j.1742-4658.2012.08609.x

6 Huang, K.-L. et al. Integrator Recruits Protein Phosphatase 2A to Prevent Pause Release and Facilitate Transcription Termination. Molecular Cell 80, 345–358.e349 (2020). 10.1016/j.molcel.2020.08.016

7 Zheng, H. et al. Identification of Integrator-PP2A complex (INTAC), an RNA polymerase II phosphatase. Science 370, eabb5872 (2020). doi:10.1126/science.abb5872

8 Fianu, I. et al. Structural basis of Integrator-mediated transcription regulation. Science 374, 883–887 (2021). doi:10.1126/science.abk0154

9 Vervoort, S. J. et al. The PP2A-Integrator-CDK9 axis fine-tunes transcription and can be targeted therapeutically in cancer. Cell 184, 3143–3162.e3132 (2021). 10.1016/j.cell.2021.04.022

10 Sangodkar, J. et al. All roads lead to PP2A: exploiting the therapeutic potential of this phosphatase. The FEBS journal 283, 1004–1024 (2016). 10.1111/febs.13573

11 Seshacharyulu, P., Pandey, P., Datta, K. & Batra, S. K. Phosphatase: PP2A structural importance, regulation and its aberrant expression in cancer. Cancer letters 335, 9–18 (2013). 10.1016/j.canlet.2013.02.036

12 Olsen, J. V. et al. Global, In Vivo, and Site-Specific Phosphorylation Dynamics in Signaling Networks. Cell 127, 635–648 (2006). 10.1016/j.cell.2006.09.026

13 Xing, Y. et al. Structure of protein phosphatase 2A core enzyme bound to tumor-inducing toxins. Cell 127, 341–353 (2006). 10.1016/j.cell.2006.09.025

14 Xu, Y. et al. Structure of the Protein Phosphatase 2A Holoenzyme. Cell 127, 1239–1251 (2006). 10.1016/j.cell.2006.11.033

15 Cho, U. S. & Xu, W. Crystal structure of a protein phosphatase 2A heterotrimeric holoenzyme. Nature 445, 53–57 (2007). 10.1038/nature05351

16 Pallas, D. C. et al. Polyoma small and middle T antigens and SV40 small t antigen form stable complexes with protein phosphatase 2A. Cell 60, 167–176 (1990). 10.1016/0092-8674(90)90726-U

17 Walter, G. & Mumby, M. Protein serine/threonine phosphatases and cell transformation. Biochimica et Biophysica Acta (BBA) - Reviews on Cancer 1155, 207–226 (1993). 10.1016/0304-419X(93)90005-W

18 Chen, Y. et al. Structural and biochemical insights into the regulation of protein phosphatase 2A by small t antigen of SV40. Nat Struct Mol Biol 14, 527–534 (2007). 10.1038/nsmb1254

19 Pim, D., Massimi, P., Dilworth, S. M. & Banks, L. Activation of the protein kinase B pathway by the HPV-16 E7 oncoprotein occurs through a mechanism involving interaction with PP2A. Oncogene 24, 7830–7838 (2005). 10.1038/sj.onc.1208935

20 Garibal, J. et al. Truncated Form of the Epstein-Barr Virus Protein EBNA-LP Protects against Caspase-Dependent Apoptosis by Inhibiting Protein Phosphatase 2A. Journal of Virology 81, 7598–7607 (2007). 10.1128/jvi.02435-06

21 Siamer, S. et al. Expression of the Bacterial Type III Effector DspA/E in Saccharomyces cerevisiae Down-regulates the Sphingolipid Biosynthetic Pathway Leading to Growth Arrest*. Journal of Biological Chemistry 289, 18466–18477 (2014). 10.1074/jbc.M114.562769

22 Jin, L. et al. Direct and Indirect Targeting of PP2A by Conserved Bacterial Type-III Effector Proteins. PLOS Pathogens 12, e1005609 (2016). 10.1371/journal.ppat.1005609

23 Li, H. et al. Pathogen protein modularity enables elaborate mimicry of a host phosphatase. Cell 186, 3196–3207.e3117 (2023). 10.1016/j.cell.2023.05.049

24 Kamoun, S. et al. The Top 10 oomycete pathogens in molecular plant pathology. Molecular Plant Pathology 16, 413–434 (2015). 10.1111/mpp.12190

25 Franceschetti, M. et al. Effectors of Filamentous Plant Pathogens: Commonalities amid Diversity. Microbiology and Molecular Biology Reviews 81, 10.1128/mmbr.00066-00016 (2017). https://doi.org/10.1128/mmbr.00066-16

26 Jiang, R. H. Y., Tripathy, S., Govers, F. & Tyler, B. M. RXLR effector reservoir in two Phytophthora species is dominated by a single rapidly evolving superfamily with more than 700 members. Proceedings of the National Academy of Sciences 105, 4874–4879 (2008). doi:10.1073/pnas.0709303105

27 He, J. et al. Structural analysis of Phytophthora suppressor of RNA silencing 2 (PSR2) reveals a conserved modular fold contributing to virulence. Proceedings of the National Academy of Sciences 116, 8054–8059 (2019). 10.1073/pnas.1819481116

28 Farkas, I., Dombrádi, V., Miskei, M., Szabados, L. & Koncz, C. Arabidopsis PPP family of serine/threonine phosphatases. Trends in plant science 12, 169–176 (2007). 10.1016/j.tplants.2007.03.003

29 Abramson, J. et al. Accurate structure prediction of biomolecular interactions with AlphaFold 3. Nature 630, 493–500 (2024). 10.1038/s41586-024-07487-w

30 Xu, Y., Chen, Y., Zhang, P., Jeffrey, P. D. & Shi, Y. Structure of a Protein Phosphatase 2A Holoenzyme: Insights into B55-Mediated Tau Dephosphorylation. Molecular Cell 31, 873–885 (2008). 10.1016/j.molcel.2008.08.006

31 Wlodarchak, N. et al. Structure of the Ca2+-dependent PP2A heterotrimer and insights into Cdc6 dephosphorylation. Cell Research 23, 931–946 (2013). 10.1038/cr.2013.77

32 Myles, T., Schmidt, K., Evans, D. R. H., Cron, P. & Hemmings, B A. Active-site mutations impairing the catalytic function of the catalytic subunit of human protein phosphatase 2A permit baculovirus-mediated overexpression in insect cells. Biochemical Journal 357, 225–232 (2001). 10.1042/bj3570225

33 Harada, K.-I. et al. Detection and identification of microcystins in the drinking water of Haimen City, China. Natural Toxins 4, 277–283 (1996). 10.1002/(SICI)(1996)4:6<277::AID-NT5>3.0.CO;2-1

34 Ueno, Y. et al. Detection of microcystins, a blue-green algal hepatotoxin, in drinking water sampled in Haimen and Fusui, endemic areas of primary liver cancer in China, by highly sensitive immunoassay. Carcinogenesis 17, 1317–1321 (1996). 10.1093/carcin/17.6.1317

35 Hou, Y. et al. A Phytophthora Effector Suppresses Trans-Kingdom RNAi to Promote Disease Susceptibility. Cell host & microbe 25, 153–165 e155 (2019). 10.1016/j.chom.2018.11.007

36 Dong, S. & Ma, W. How to win a tug-of-war: the adaptive evolution of Phytophthora effectors. Current opinion in plant biology 62, 102027 (2021). 10.1016/j.pbi.2021.102027

37 Ma, W. & Guttman, D. S. Evolution of prokaryotic and eukaryotic virulence effectors. Current opinion in plant biology 11, 412–419 (2008). 10.1016/j.pbi.2008.05.001

38 Bialojan, C. & Takai, A. Inhibitory Effect of a Marine-Sponge Toxin, Okadaic Acid, on Protein Phosphatases - Specificity and Kinetics. Biochem J 256, 283–290 (1988). DOI 10.1042/bj2560283

39 Wang, Y., Bouwmeester, K., Van de Mortel, J. E., Shan, W. X. & Govers, F. A novel Arabidopsis-oomycete pathosystem: differential interactions with reveal a role for camalexin, indole glucosinolates and salicylic acid in defence. Plant Cell Environ 36, 1192–1203 (2013). 10.1111/pce.12052

40 Schorb, M., Haberbosch, I., Hagen, W. J. H., Schwab, Y. & Mastronarde, D. N. Software tools for automated transmission electron microscopy. Nature Methods 16, 471–477 (2019). 10.1038/s41592-019-0396-9

41 Mastronarde, D. N. Automated electron microscope tomography using robust prediction of specimen movements. Journal of Structural Biology 152, 36–51 (2005). 10.1016/j.jsb.2005.07.007

42 Punjani, A., Rubinstein, J. L., Fleet, D. J. & Brubaker, M. A. cryoSPARC: algorithms for rapid unsupervised cryo-EM structure determination. Nature Methods 14, 290–296 (2017). 10.1038/nmeth.4169

43 Emsley, P., Lohkamp, B., Scott, W. G. & Cowtan, K. Features and development of Coot. Acta Crystallographica. Section D, Biological Crystallography 66, 486–501 (2010). 10.1107/S0907444910007493

44 Adams, P. D. et al. PHENIX: a comprehensive Python-based system for macromolecular structure solution. Acta Crystallographica Section D 66, 213–221 (2010). doi:10.1107/S0907444909052925

45 Pettersen, E. F. et al. UCSF ChimeraX: Structure visualization for researchers, educators, and developers. Protein Science 30, 70–82 (2021). 10.1002/pro.3943

