## supplemental Figures and table for "Pathogen effector forms a hexameric phosphatase holoenzyme with host core enzyme to promote disease"

**This file contains:**

Supplementary Fig. 1-8

Supplementary Table 1

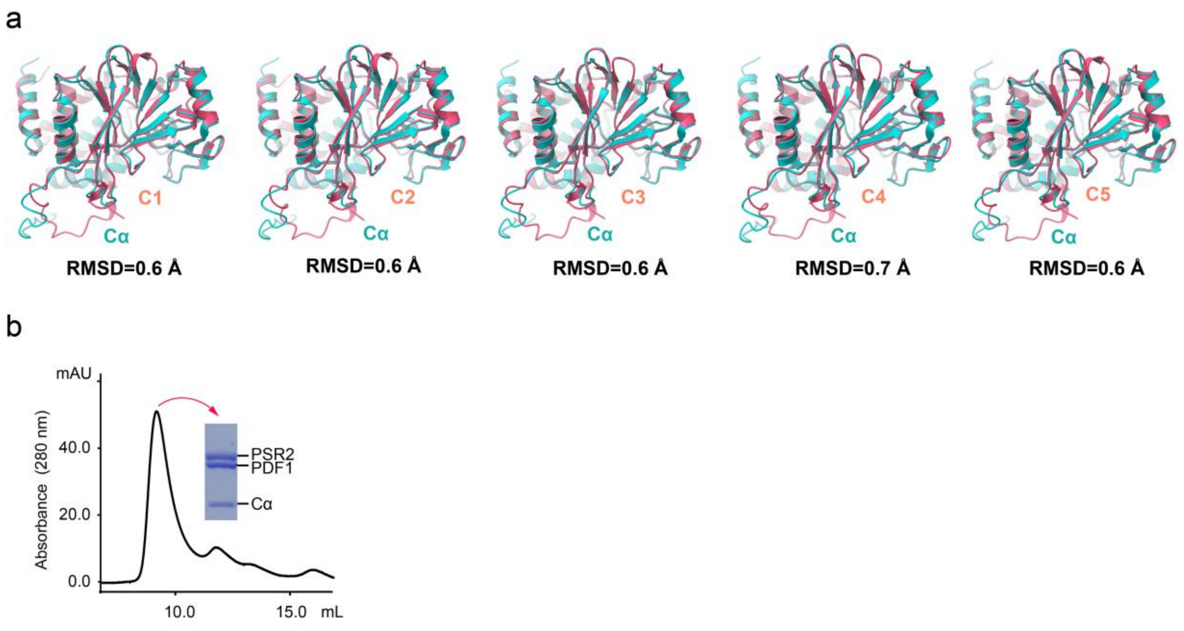

**Supplementary Fig.1 The human PP2A C subunit Cα is suitable to reconstitute PSR2-PDF1-Cα complex.**

**a**. Structural comparison between human PP2A C subunit Cα from PDB code 2iae and predicted structures of the Arabidopsis PP2A C subunits (predicted by alpha fold 3) shows that their structures are almost the same.

**b**. Gel filtration chromatography of PSR2-PDF1-Cα used for cryo-EM sample preparation.

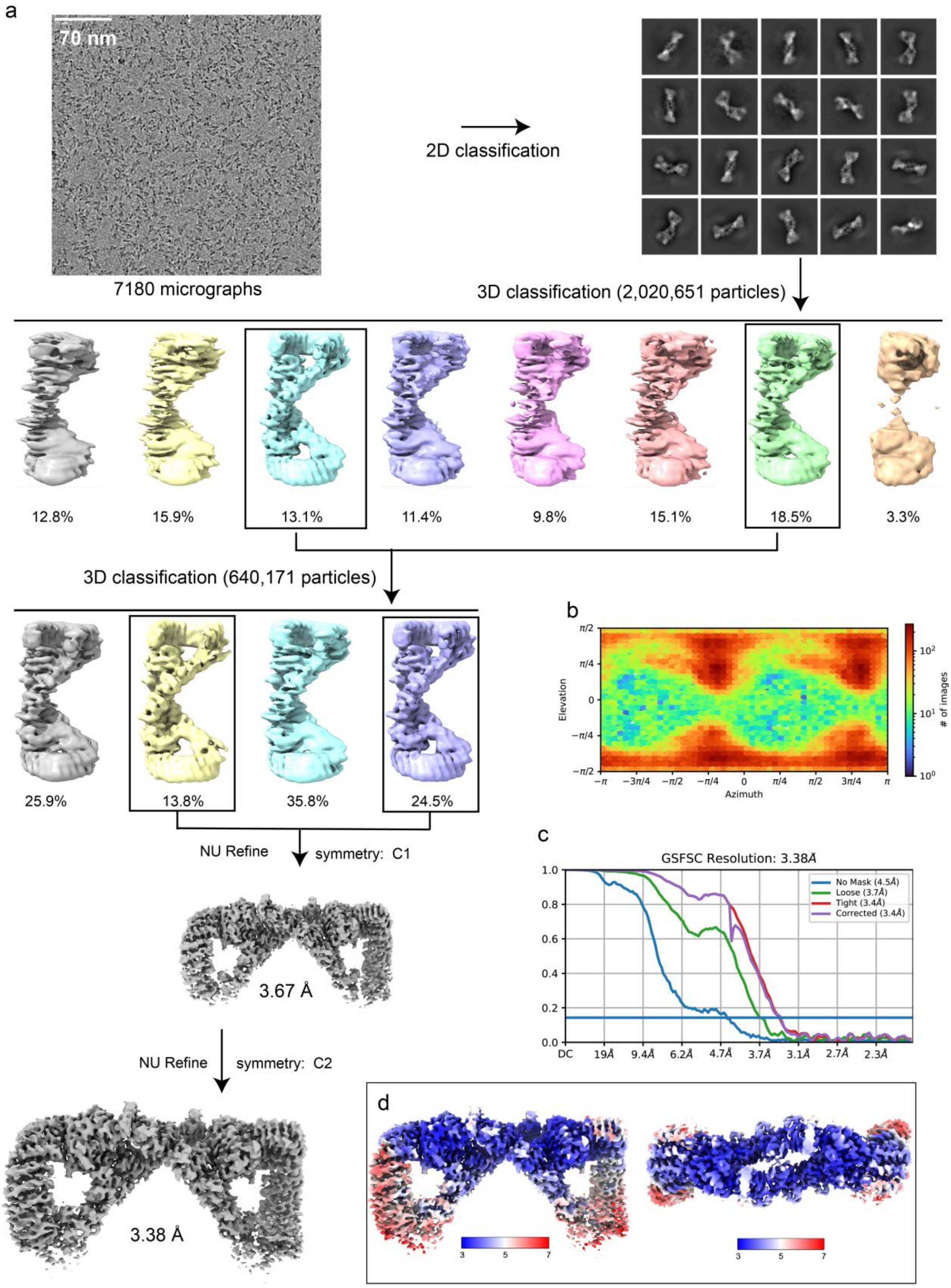

**Supplementary Fig.2 The processed result of PSR2$^{hexamer}$-PDF1-Cα Cryo-EM dataset.**

**a**. the cryo-EM data processed workflow of PSR2$^{hexamer}$-PDF1-Cα.

**b**. Angular distribution of the $PSR2^{hexamer}$-PDF1-Cα particles.

**c**. Fourier shell correlation (FSC) curve.

**d**. Local resolutions distribution of the $PSR2^{hexamer}$-PDF1-Cα cryo-EM map.

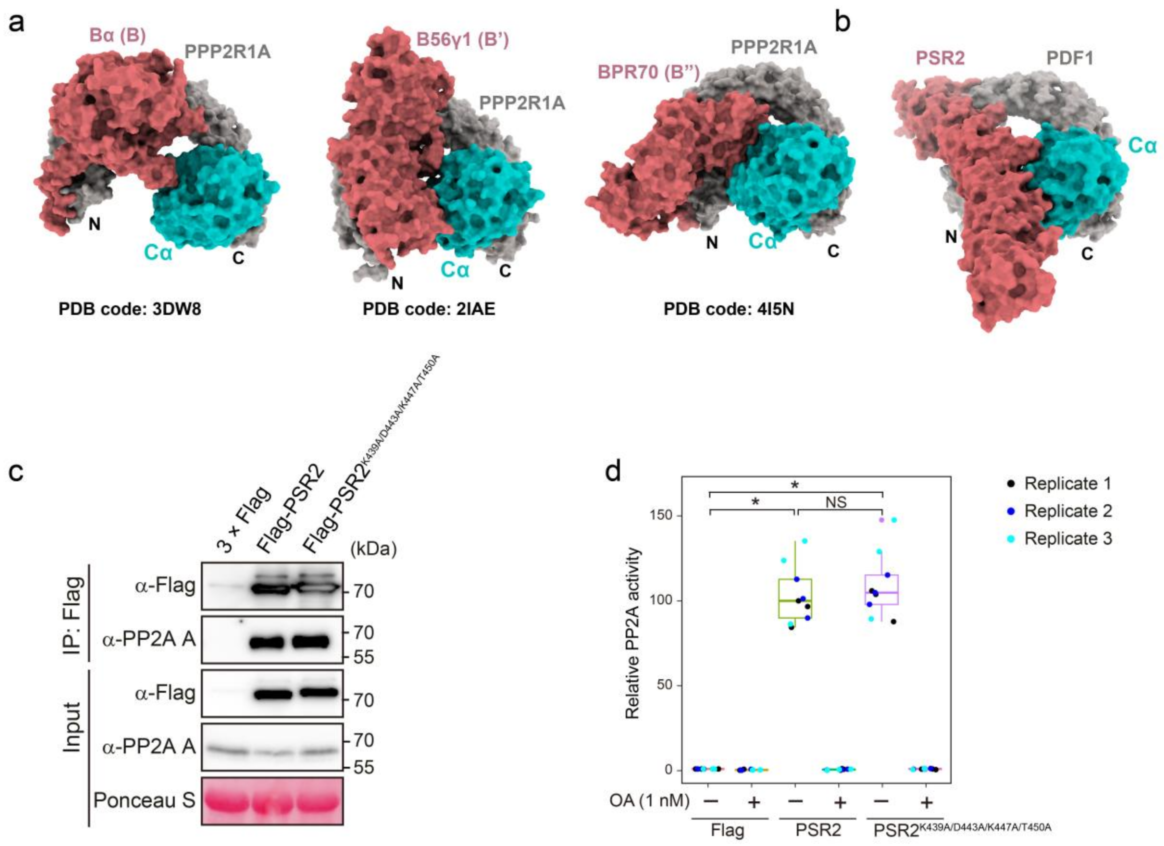

**Supplementary Fig.3 Structural comparison of different PP2A holoenzymes, and experiments show that the module formed by LWY4-LWY5 does not exhibit a higher importance for PSR2-PDF1-Cα hexamer holoenzyme formation.**

**a-b**. Structural comparison between endogenous holoenzymes (A) and heterotrimeric holoenzyme (B) separated from PSR2-PDF1-Cα hexameric holoenzyme.

**c**. PSR2$^{K439A/D443A/K447A/T450A}$ mutant does not influence its interaction with PP2A core enzyme *in planta* FLAG-tagged PSR2 or PSR2$^{K439A/D443A/K447A/T450A}$ were expressed in *Nicotiana benthamiana* and immunoprecipitated using anti-FLAG magnetic beads.

**d**. PSR2$^{K439A/D443A/K447A/T450A}$ mutant does not influence its phosphatase activity. Protein complexes enriched by using an anti-PSR2 antibody were examined for phosphatase activity using a phosphopeptide as the substrate. Okadaic acid (OA) is a PP2A specific inhibitor.

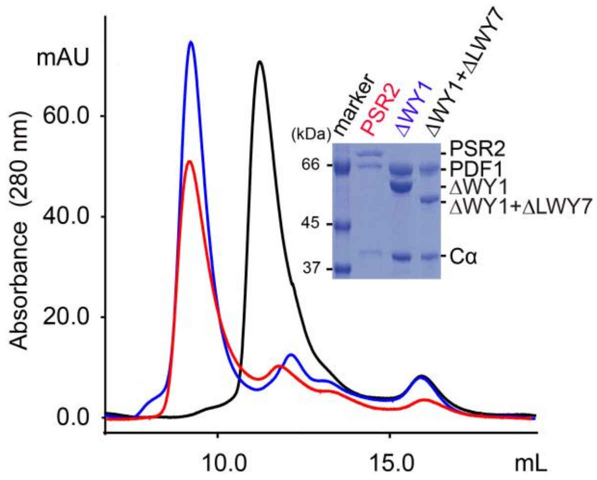

**Supplementary Fig.4 Gel filtration chromatography results show that PSR2$^{\Delta WY1}$-PDF1-Cα can still exist as a hexamer but exist as a heterotrimer after truncating the WY1 and LWY7 unit.**

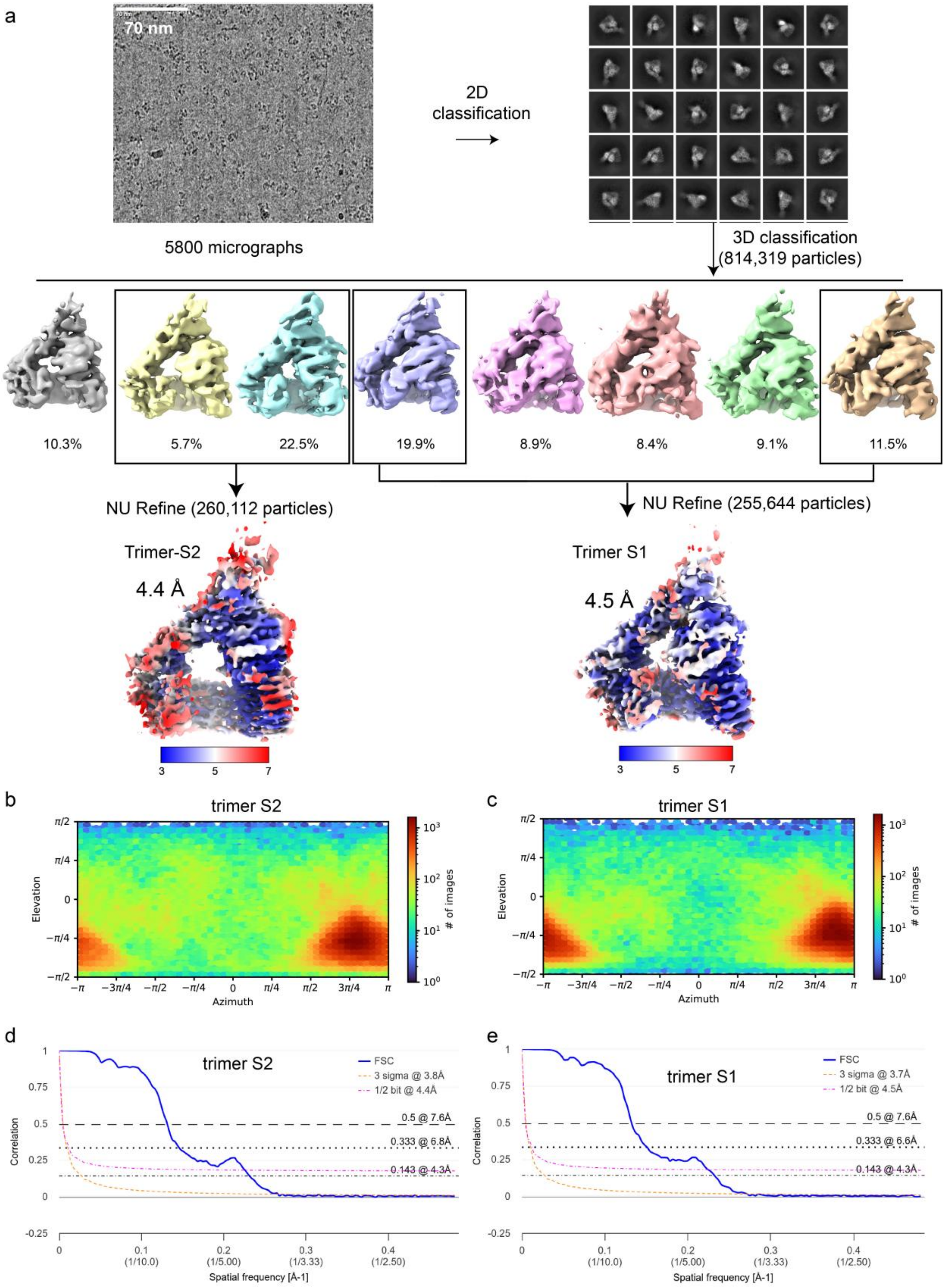

**Supplementary Fig.5 The processed result of PSR2$^{heterotrimer}$-PDF1-Cα cryo-EM dataset.**

**a**. the cryo-EM data processed workflow of PSR2$^{heterotrimer}$-PDF1-Cα.

**b-c**. Angular distribution of the PSR2$^{heterotrimer}$-PDF1-Cα particles in Trimer-S2 (**b**) and Trimer-S1 (**c**), respectively.

**d-e**. FSC curve of the PSR2$^{heterotrimer}$-PDF1-Cα particles in Trimer-S2 (**d**) and Trimer-S1 (**e**), respectively. The evaluated resolution by Cryosparc is not accurate, so, we chose the accurate resolution evaluation data analysed by wwPDB Validation System (https://validate-rcsb-1.wwpdb.org/).

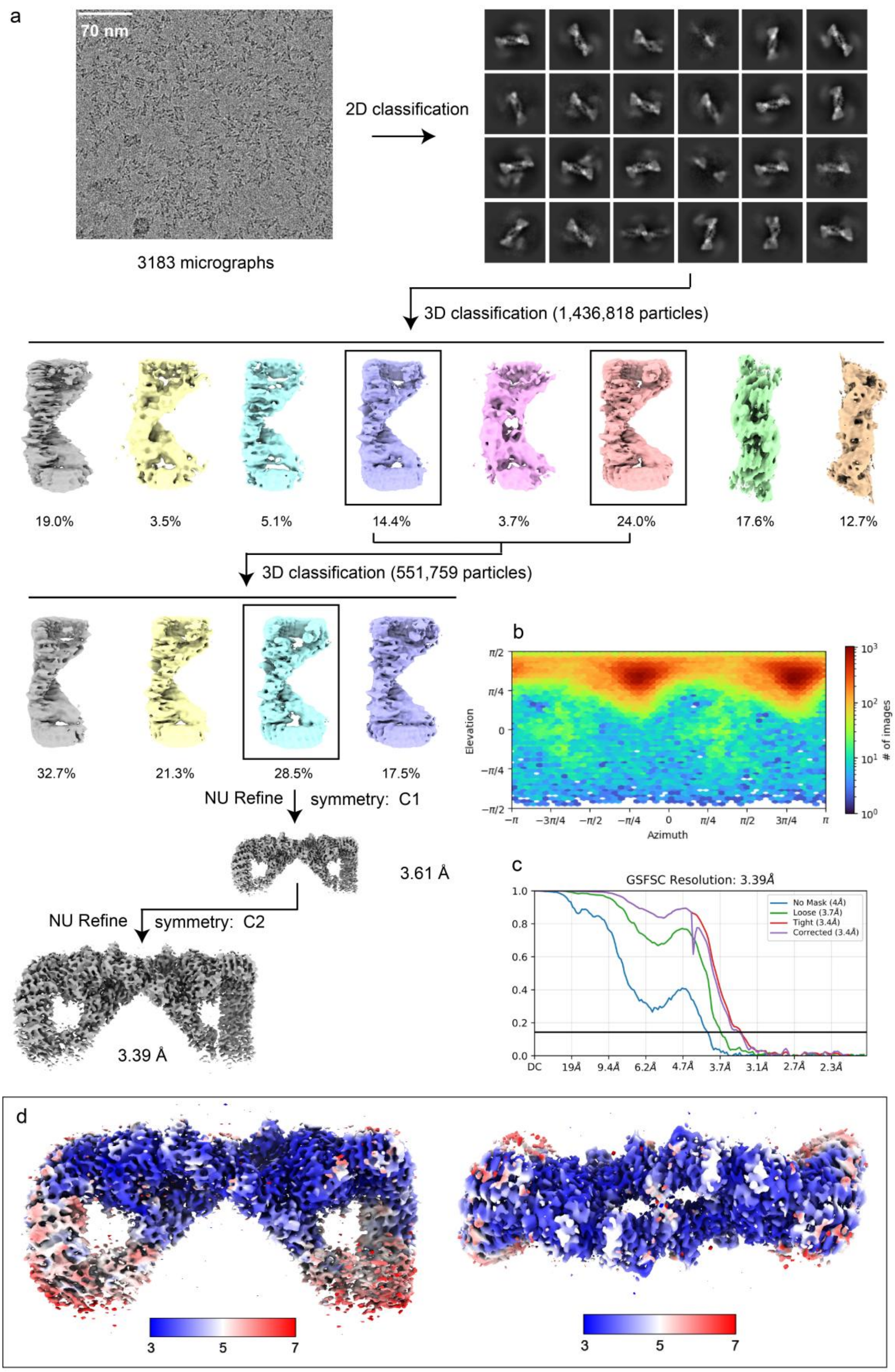

**Supplementary Fig.6 The processed result of PSR2$^{hexamer}$-PDF1-Cα$^{substrate}$ (substrate bound hexamer) Cryo-EM dataset.**

**a**. The cryo-EM data processed workflow of PSR2 $^{hexamer}$-PDF1-Cα$^{substrate}$.

**b**. Angular distribution of the PSR2 $^{hexamer}$-PDF1-Cα$^{substrate}$ particles.

**c**. FSC curve.

**d**. local resolutions distribution of the PSR2 $^{hexamer}$-PDF1-Cα$^{substrate}$ cryo-EM map.

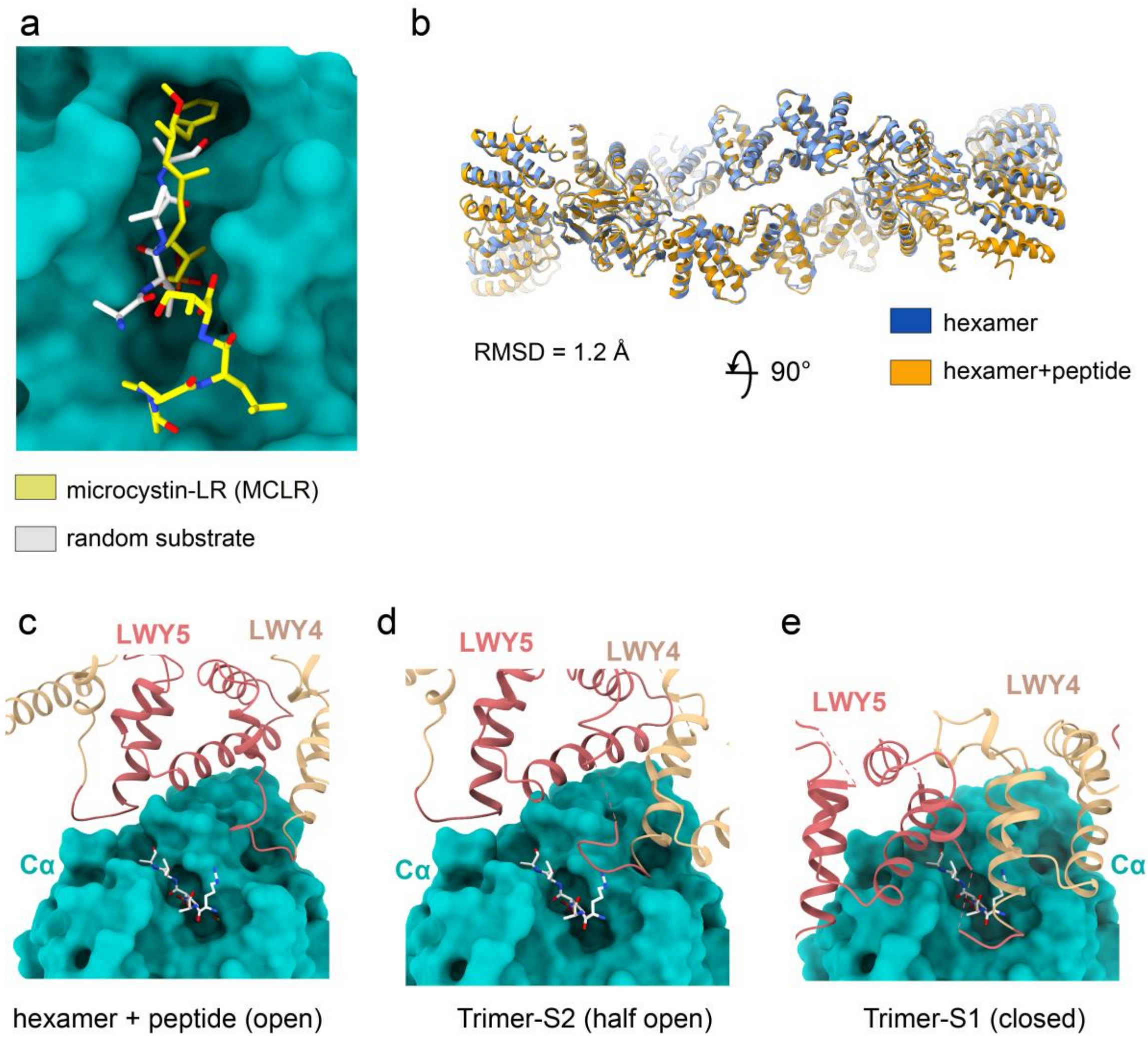

**Supplementary Fig.7 Comparison of structural differences between substrate-bound and substrate-unbound states.**

**a**. Superposition of C subunits from PSR2$^{hexamer}$-PDF1-Cα$^{substrate}$ and endogenous holoenzyme (PDB code: 2IAE) shows that the binding sites of random substrate and specific inhibitor microcystin-LR are overlapped.

**b**. Overall structural superposition of PSR2$^{hexamer}$-PDF1-Cα and PSR2 $^{hexamer}$-PDF1-Cα$^{substrate}$ shows that PSR2$^{hexamer}$-PDF1-Cα does not have obvious conformational change after binding to random substrate.

**c**. The relative position of random substrate and LWY4-LWY5 in PSR2$^{hexamer}$-PDF1-Cα$^{substrate}$ shows that PSR2 stay away from the random substrate and does not influence the access of substrate to the catalytic pocket.

**d**. Superposition of C subunits from PSR2$^{hexamer}$-PDF1-Cα$^{substrate}$ and Trimer-S2 shows that the

PSR2 in Trimer-S2 stays close to and does not cover the substrate, indicating in Trimer-S2 PSR2 probably does not highly affect the access of substrate to the catalytic pocket.

**e**. Superposition of C subunits from $PSR2^{hexamer}$-PDF1-$C\alpha^{substrate}$ and Trimer-S1 shows that the PSR2 in Trimer-S1 and the random substrate are overlapped, indicating in Trimer-S1, PSR2 probably affect the access of substrate to the catalytic pocket.

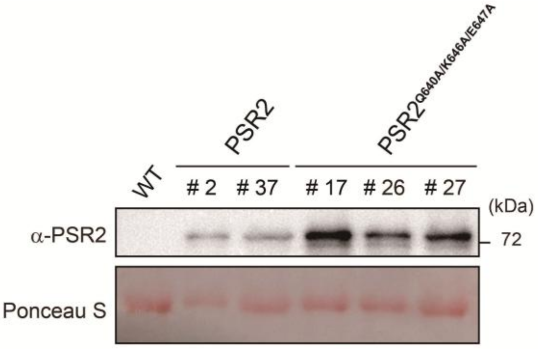

**Supplementary Fig.8 The expression level of PSR2 and PSR2$^{Q640A/K646A/E647A}$ in the transgenic plants.**

Western blotting showing the protein levels of PSR2 and PSR2$^{Q640A/K646A/E647A}$ in the constitutive transgenic Arabidopsis plants. Total proteins were extracted from 2-week-old Arabidopsis seedlings and PSR2 and PSR2$^{Q640A/K646A/E647A}$ was detected using an anti-PSR2 antibody. Ponceau S staining was used to confirm equal loading.

**Supplementary Table 1. Cryo-EM data collection, refinement and validation statistics**

| | PDF1-Cα-PSR2 (EMDB-61142) (PDB code : 9J5I) | PDF1-Cα$^{substrate}$-PSR2 (EMDB-61144) (PDB code : 9J5K) | trimer-S2 (EMDB-61147) (PDB code : 9J5N) | Trimer-S1 (EMDB-61150) (PDB code : 9J5R) |
|---|---|---|---|---|
| **Data collection and processing** | | | | |
| Magnification | 130,000 | 130,000 | 130,000 | 130,000 |
| Voltage (kV) | 300 | 300 | 300 | 300 |
| Electron exposure (e–/Å$^2$) | 50 | 50 | 50 | 50 |
| Defocus range (μm) | 1.2-1.8 | 1.2-1.8 | 1.2-1.8 | 1.2-1.8 |
| Pixel size (Å) | 1.04 | 1.04 | 1.04 | 1.04 |
| Symmetry imposed | C2 | C2 | C1 | C1 |
| Initial particle images (no.) | 7180 | 3183 | 5800 | 5800 |
| Final particle images (no.) | 6807 | 3012 | 5273 | 5273 |
| Map resolution (Å) | 3.38 | 3.27 | 4.4 | 4.5 |
| FSC threshold | 0.143 | 0.143 | 0.143 | 0.143 |
| **Refinement** | | | | |
| Initial model used (PDB code) | 7XVK | 7XVK | 7XVK | 7XVK |
| Map sharpening *B* factor (Å$^2$) | -80.5 | -85.1 | -126.1 | -93.0 |
| Model composition | | | | |
| Non-hydrogen atoms | 19306 | 19058 | 7549 | 8322 |
| Protein residues | 2698 | 2702 | 1077 | 1171 |
| $Mn^{2+}$ | 4 | 4 | 2 | 2 |
| *B* factors (Å$^2$) | | | | |
| Protein | 113.24 | 87.59 | 75.33 | 85.03 |
| $Mn^{2+}$ | 69.20 | 38.89 | 44.53 | 50.44 |
| R.m.s. deviations | | | | |
| Bond lengths (Å) | 0.003 | 0.002 | 0.002 | 0.003 |
| Bond angles (°) | 0.755 | 0.566 | 0.559 | 0.710 |
| Validation | | | | |
| MolProbity score | 2.22 | 2.04 | 2.22 | 2.05 |
| Clashscore | 8.92 | 9.92 | 9.09 | 12.08 |
| Ramachandran plot | | | | |
| Favored (%) | 94.81 | 95.44 | 94.36 | 93.44 |
| Allowed (%) | 5.19 | 4.56 | 5.64 | 6.56 |
| Disallowed (%) | 0.00 | 0.00 | 0.00 | 0.00 |
